# Ultrafast metaproteomics for quantitative assessment of strain isolates and microbiomes

**DOI:** 10.1101/2024.04.15.589175

**Authors:** Elizaveta Kazakova, Mark Ivanov, Tomiris Kusainova, Julia Bubis, Valentina Polivtseva, Kirill Petrikov, Vladimir Gorshkov, Frank Kjeldsen, Mikhail Gorshkov, Yanina Delegan, Inna Solyanikova, Irina Tarasova

## Abstract

**BACKGROUND:** Microbial communities play a crucial role in human health and environmental regulation, but present an especial challenge for the analytical science due to their diversity and dynamic range. Tandem mass spectrometry provides functional insights on microorganisms life cycle, but still lacks throughput and sensitivity. MALDI TOF is widely used for ultrafast identification of species, but does not assess their functional activity. Development of ultrafast mass spectrometry methods and bioinformatic approaches applicable for both accurate identification and functional assessment of microbial communities based on their protein content is of high interest.

**RESULTS:** We show for the first time that both identity and functional activity of microorganisms and their communities can be accurately determined in experiments as short as 7 minutes per sample, using the basic Orbitrap MS configuration without peptide fragmentation. The approach was validated using strain isolates, mock microbiomes composed of bacteria spiked at known concentrations and human fecal microbiomes. Our new bioinformatic algorithm identifies the bacterial species with an accuracy of 95 %, when no prior information on the sample is available. Microbiome composition was resolved at the genus level with the mean difference between the actual and identified components of 12 %. For mock microbiomes, Pearson coefficient of up to 0.97 was achieved in estimates of strain biomass change. By the example of *Rhodococcus* biodegradation of *n*-alkanes, phenols and its derivatives, we showed the accurate assessment of functional activity of strain isolates, compared with the standard label-free and label-based approaches.

**SIGNIFICANCE:** Our approach makes microbial proteomics fast, functional and insightful using the Orbitrap instruments even without employing peptide fragmentation technology. The approach can be applied to any microorganisms and can take a niche in routine functional assessment of microbial pathogens and consortiums in clinical diagnostics together with MALDI TOF MS and 16S rRNA gene sequencing.

## Introduction

Metaproteomics is a growing area of research^1^ devoted to monitoring the physiology of biological communities, for example, microbial ones, at the level of their proteomes^2^. The metaproteomic studies are focused on determining the species diversity in a community, studying various stages of the life cycle of microorganisms, their response to stress, the functional characterization of microbiomes, etc^3^. Being a young field, metaproteomics possess challenges that must be overcome before using for routine research^2,4^. Currently, the research efforts are focused on accuracy, sensitivity, and reproducibility of the analysis, development of bioinformatic solutions, and standardization of workflows^5^.

Proteomic analysis of microbiomes relies on liquid chromatography tandem mass spectrometry (LC-MS/MS) that allows the identification of thousands of protein molecules in a single experiment. A common approach is data-dependent acquisition (DDA) mass spectrometry, in which the success of peptide identification strongly depends on the choice of the most abundant ions. Reportedly, the performance of DDA decreases with increasing the sample complexity^6^. The time-consuming DDA analysis with insufficient peptide fragmentation and protein sequence coverage, and heavy bioinformatic processing of metaproteomic data still present a gap for more efficient solutions. Another acquisition method recently applied in metaproteomics is data independent acquisition (DIA)^7,8,9^. DIA allows collecting fragment spectra for all peptide ions without pre-selection of a fixed number of top abundant ones, and, thus, promises better sequence coverage and protein quantitation^10^. The third approach can be the use of short chromatographic gradients with acquisition of only MS1 spectra without fragmentation. It has been shown that interpretation of LC-MS1 data based on alignments of peptide feature intensities without peptide and protein identification allows large-scale screening of microbiomes and characterization of changes in MS1 profiles, depending on drug treatment^11^. Widely implemented in clinics, MALDI-TOF MS allows identification of the genus and species by matching mass spectra against spectral libraries of known microorganisms^12^, but the strain resolution with this technique is problematic^13^. Identifying bacteria by matching the experimental LC-MS1 data against *in silico* generated taxon-specific tryptic peptide masses compiled based on UniProt, SwissProt, and TrEMBL databases, can resolve species and, sometimes, strain identity^14^. However, the functional characterization of microbiomes still cannot be assessed without the identification of proteins.

Identification of proteins relies on protein databases which represents one of the hardest issues in MS-based metaproteomics due to its huge size. The choice of the protein database affects the sensitivity of peptide identification because the larger the search space, the higher the chance of random and incorrect matches. The optimal database should only contain proteins that are present in the sample^15^. From this viewpoint, the protein database constructed using metagenome data or restricted using 16S rRNA gene analysis is the best. However, they are costly and require large sample amounts, as it is important to obtain high-quality sequences of the genome. Attempts to fix the FDR problem typically include selecting top organisms or peptide sequences within a certain window of *m/z* values followed by repeating the search using the restricted database^16,17,18,19,20,21^. The MetaProteomeAnalyzer^22^ implements a search with four different search engines and the integration of the results. Following the search, the identified proteins are grouped into so-called meta-proteins and annotated with protein-level information.

This study presents an adaptation of the novel ultrafast MS1-based method called DirectMS1^23,24^ to the characterization of single microorganisms and microbiomes. Using DirectMS1 we demonstrate the blind identification (with no prior knowledge of the bacteria in the sample) of a single bacterial species and microbial compositions using two-step database search against the microbial part of SwissProt and TrEMBL (62Gb database size), the evaluation of fold changes in biomass of microorganisms between the metaproteomic samples, and the assessment of metabolic activity of *Rhodococcus* species responding to a change in growth conditions and cold shock.

## Methods

### Public datasets

Public dataset (https://zenodo.org/record/3573994) collected in DDA mode for 19 well-characterized bacterial strains^14^ was used to validate the two-step search algorithm for blind identification of individual microorganisms against the microbial part of SwissProt and TrEMBL database (accessed on Jan 2020).

Public dataset (PRJEB42466) for human fecal microbiome, collected in DDA mode, was used for the testing of the blind search algorithm to identify composition of human fecal microbiome^5^.

### Strains, genomes and proteogenomic databases

Strains used in the study were isolated from different sources and characterized as previously described (*Rhodococcus erythropolis* X5^25^, *Rhodococcus opacus* S8^26^, *Rhodococcus qingshengii* 7B^27^, *Rhodococcus qingshengii* VT6^28^, *Priestia aryabhattai* 25^29^, *Gordonia amicalis* 6-1^30^, *Gordonia alkanivorans* 135^31,32^, *Gordonia polyisoprenivorans* 135^33^, *Rhodococcus opacus* 1CP^34^, *Rhodococcus opacus* 3D^35^).

To construct protein databases using genome data, NCBI PGAP 6.6^36^ was used. Annotations of proteins of interest were manually checked using UniProt^37^.

### Cell cultivation to characterize metabolic activity changes in response to stress

*R. opacus* 1CP was cultivated in the presence of four different carbon sources (glucose, benzoate, phenol, 4-chlorophenol) with six biological replicates per condition. The cultivation was carried out in 750-mL Erlenmeyer flasks with 200 mL of the mineral medium of the following composition (g/L): Na_2_HPO_4_ - 0.7, K_2_HPO_4_ - 0.5, NH_4_NO_3_ - 0.75, MgSO_4_×7H_2_O - 0.2, MnSO_4_ - 0.001, FeSO_4_ - 0.02 with the addition of one of the following sources of carbon: phenol 500 mg/L, benzoate 500 mg/L, 4-chlorophenol 100 mg/L, glucose 10 g/L. The cultivation was performed for 48-72 hours in a shaking incubator (HZQ-111) at 24℃ and 220 rpm. During cultivation time, the carbon sources (phenol and 4-chlorophenol) were periodically injected into the medium after consumption of previously added. The disappearance of substrates was determined by characteristic absorption spectra in the 230-290 nm region. Culture growth was monitored with an optical absorption coefficient at 560 nm.

*R. erythropolis* X5 cells were grown in an orbital shaking incubator at 180 rpm at two different temperatures (28°C and 6°C) for 3 and 11 days, respectively, with three biological replicates. The cultivation was carried out in 750 mL Erlenmeyer flasks with 100 mL of the modified Evans mineral salts medium supplemented with 2 mL of *n*-hexadecane. The composition of the cultivation medium was as follows (per liter): K_2_HPO_4_, 8.71 g; 5 M solution of NH_4_Cl, 1 mL; 0.1 M solution of Na_2_SO_4_, 1 mL; 62 mM solution of MgCl_2_, 1 mL; 1 mM solution of CaCl_2_, 1 mL; 0.005 mM solution of (NH_4_)_6_Mo_7_O_24_, 1 mL; trace element solution, 1 mL. The value of рН was adjusted to 7.5 by concentrated HCl. Composition of the trace element solution in 1 % water solution of HCl (g/L): ZnO, 0.41 g; FeCl_3_, 2.9 g; MnCl_2_, 1.28 g; CuCl_2_, 0.13 g; CoCl_2_, 0.26 g; H_3_BO_3_, 0.06 g.

### Cell cultivation for model microbiomes

Cells were cultivated for 72 hours in a shaking incubator (HZQ-111) at 27°С and 250 rpm in Erlenmeyer flasks with 100 mL of Evans medium. The composition of the medium was as follows (g/L or mL/L:): K_2_HPO_4_ 8.71 g, 5 M NH_4_CI 1 mL, 0.1 M Na_2_SO_4_ 1 mL, 62 mM MgCI_2_ 1 mL, 1 mM CaCI_2_ 1 mL, 0.005 mM (NH_4_)_6_Mo_7_O_24_ × 4H_2_O, micronutrients solution 1 mL (as follows in g/L): ZnO 0.41 g, FeCI_3_ × 6H_2_O 5.4 g, MnCI_2_ × 4H_2_O 2 g, CuCI_2_ × 2H_2_O 0.17 g, CoCI_2_ × 6H_2_O 0.48 g, H_3_BO_3_ 0.06 g, (pH 7.0) - with the addition of 200 uL 10^8^ CFU inoculum and glucose (1 % v/v for *R. opacus* 1CP and 2 % v/v for *R. erythropolis* X5, *G. alkanivorans* 135, *G. amicalis*, *P. aryabhattai* 25) as the sole source of carbon and energy.

### Model microbiome compositions

Model microbiomes were prepared by mixing tryptic peptide solutions to test the performance of DirectMS1 method. The composition of each sample is described in **Tables 1–3**.

**Table 1.** Model I: five low fold-change mixtures of peptides (ng) from 3 to 5 strains, total of 500 ng of the sample per injection.

| Strain | M1.1 | M1.2 | M1.3 | M1.4 | M1.5 |
| --- | --- | --- | --- | --- | --- |
| *Rhodococcus opacus* 1CP | 167 | 250 | 167 | 100 | 71 |
| *Rhodococcus erythropolis* X5 | - | - | - | 100 | 143 |
| *Priestia aryabhattai* 25 | 167 | 125 | 250 | 100 | 71 |
| *Gordonia amicalis* 6-1 | - | - | - | 100 | 143 |
| *Gordonia alkanivorans* 135 | 167 | 125 | 83 | 100 | 71 |

### Sample preparation for mass spectrometry-based proteomics

For mix models I and III (Tables 1 and 3), cell pellets were resuspended in 150 µL lysis buffer (50 mM ammonium bicarbonate (ABC), 10 % ACN, 0.1 % ProteaseMax (Promega, USA)). Cells incubated for 45 min at room temperature, after that they were boiled for 10 min at 95 °C, and then cooled down on ice. To lyse cells, the samples were sonicated for 10 min (1 s on 1 s off) on 60 % amplitude (QSonica Q125, Newtown, Connecticut, USA). Samples were centrifuged for 7 min on 10 000 g, then supernatant was taken, and protein concentration was measured by BCA kit (Thermo Scientific, Germany). 10 mM dithiothreitol (DTT) was added for 20 min at 56 °C, followed by incubation with iodoacetamide (IAA) for 30 min in the dark at room temperature. Lys-c was added in 1:100 m/m ratio and samples incubated for 2h at 37 °C then trypsin was added in 1:50 m/m ratio and samples incubated overnight at 37 °C. Digest was stopped by adding 1 % trifluoroacetic acid (TFA), then samples were desalted with OASIS HLB cartridges (Waters, USA) and dried in a vacuum concentrator.

For mix model II (Table 2) and characterization of metabolic activity changes, cell pellets were resuspended in a 150 µL lysis buffer (100 mM ABC, 4 % SDS, 10 mM DTT). Cells were boiled for 10 min at 95°C and then cooled down on ice. To lyse cells, the samples were sonicated for 10 min (1 s on 1 s off) on 60 % amplitude (QSonica Q125, Newtown, Connecticut, USA). Samples were centrifuged for 10 min on 10 000 g, then the supernatant was taken. 100 uL of supernatant was purified and concentrated with chloroform-methanol precipitation. Dry pellets were resuspended in 50 uL of 4 M urea in 50 mM ABC, urea concentration was further reduced to 1 M with 50 mM ABC. Protein concentration was measured by a BCA kit (Thermo Scientific, Germany). 10 mM DTT was added for 20 min at 56 °C, followed by incubation with IAA for 30 min in the dark at room temperature. Trypsin was added in a 1:50 m/m ratio and samples were incubated overnight at 37 °C. Digest was stopped by adding 1 % TFA, then samples were desalted with OASIS HLB cartridges (Waters, USA) and dried in a vacuum concentrator.

**Table 2.** Model II: three mixtures of peptides (ng) from 8 strains, total of 1 ug of the sample per injection.

| Strain | M2.1 | M2.2 | M2.3 |
| --- | --- | --- | --- |
| *Rhodococcus opacus* 1CP | 16 | 125 | 16 |
| *Rhodococcus opacus* 3D | 63 | 8 | 31 |
| *Rhodococcus opacus* S8 | 251 | 63 | 4 |
| *Rhodococcus qingshengii* 7B | 31 | 502 | 251 |
| *Rhodococcus qingshengii* VT6 | 125 | 251 | 502 |
| *Rhodococcus erythropolis* X5 | 4 | 16 | 125 |
| *Gordonia alkanivorans* 135 | 8 | 4 | 63 |
| *Gordonia polyisoprenivorans* 135 | 502 | 31 | 8 |

**Table 3.**
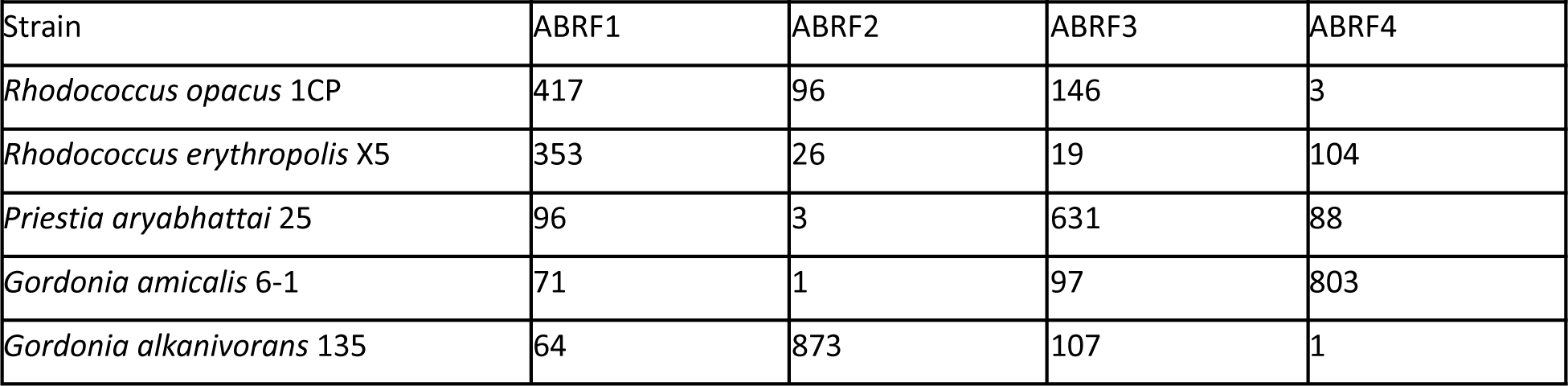
Model III: four ABRF-like mixtures of peptides (ng) from five strains, total of 1 ug of the sample per injection. Benchmark design was suggested earlier. ^38^.

| Strain | ABRF1 | ABRF2 | ABRF3 | ABRF4 |
| --- | --- | --- | --- | --- |
| *Rhodococcus opacus* 1CP | 417 | 96 | 146 | 3 |
| *Rhodococcus erythropolis* X5 | 353 | 26 | 19 | 104 |
| *Priestia aryabhattai* 25 | 96 | 3 | 631 | 88 |
| *Gordonia amicalis* 6-1 | 71 | 1 | 97 | 803 |
| *Gordonia alkanivorans* 135 | 64 | 873 | 107 | 1 |

### LC-MS1 data acquisition

The LC-MS experiments were performed using Orbitrap Q Exactive HF-X mass spectrometer (Thermo Scientific, San Jose, CA, USA) coupled to UltiMate 3000 LC system (Thermo Fisher Scientific, Germering, Germany) and Orbitrap Fusion Lumos mass spectrometer (Thermo Scientific, San Jose, CA, USA) coupled to UltiMate 3000 LC system (Thermo Fisher Scientific, Germering, Germany) and equipped with FAIMS Pro interface. Trap column μ-Precolumn C18 PepMap100 (5 μm, 300 μm, i.d. 5 mm, 100 Å) (Thermo Fisher Scientific, USA) and self-packed analytical column (Reprosil-Pur 3 μm, 75 μm i.d., 5 cm length) were employed for separation. Mobile phases were as follows: (A) 0.1 % formic acid (FA) in water; (B) 80 % ACN, 0.1 % FA in water. The gradient was from 5 % to 35 % phase B in 4.8 min at 1500 nL/min. Total method time including column washing and equilibration was 7.5 min. Field asymmetric ion mobility spectrometry (FAIMS) separations were performed with the following compensation voltages (CV) −50 V, −65 V, and −80 V in a stepwise mode during LC-MS analysis. Data acquisition was performed in MS1-only mode. Full MS scans were acquired in a range from *m*/*z* 375 to 1500 at a resolution of 120,000 at *m*/*z* 200 with AGC target of 4e5, 1 microscan, and 50 ms maximum injection time. Samples were resuspended in phase A and quantities of 1 μg were loaded per injection.

### LC-MS2, data dependent acquisition

MS/MS-based analysis of samples was performed using Orbitrap Lumos Fusion mass spectrometer (Thermo Fisher Scientific, San Jose, CA, USA) and Orbitrap Q Exactive HF mass spectrometer (Thermo Fisher Scientific, San Jose, CA, USA) coupled to UltiMate 3000 LC system. Trap column µ-Precolumn C18 PepMap100 (5 µm, 300 µm, i.d. 5 mm, 100 Å) (Thermo Fisher Scientific, USA) and self-packed analytical column (Inertsil 2 µm, 75 µm i.d., 25 cm length) were employed for separation. Mobile phases were as follows: (A) 0.1 % FA in water; (B) 95 % ACN, 0.1 % FA in water.

For TMT-based quantitation of *R. erythropolis* X5, the gradient from 5 % to 30 % phase B for 114 min at a flow rate of 300 nL/min was used. Data was acquired in top20 mode. Full MS scans were acquired in a range from m/z 300 to 1400 at a resolution of 60000 at m/z 200 with AGC (Automatic Gain Control) target of 3e6, 1 microscan, and 50 ms maximum injection time. Precursor ions were isolated in a 1.4 m/z window and accumulated for a maximum of 100 ms or until the 1e5 AGC target was reached.

For label free quantitation of *R. erythropolis* X5, the gradient from 2 % to 40 % phase B for 60 min at a flow rate of 300 nL/min was used. Full MS scans were acquired in a range from m/z 300 to 1400 at a resolution of 60000 at m/z 200 with AGC target of 3e6, 1 microscan, and 50 ms maximum injection time. Precursor ions were isolated in a 1.4 m/z window and accumulated for a maximum of 50 ms or until the AGC target of 1e5 charges was reached. Precursors of charge states from 2+ to 6+ (inclusive) were scheduled for fragmentation. To save instrument time for label free DDA analysis, biological replicates were pooled and each pooled sample was measured in triplicate.

### Data processing

Peptide feature deteсtion was performed using the Biosaur2 software^39^. The proteomic search engine ms1searchpy^24^ was used for protein identification. Searches were performed against the SwissProt + TrEMBL database (accessed on 06 Dec 2020) of bacterial proteins and strain-specific databases based on annotations of bacterial genomes. **Table 4** summarizes the origin of databases used for every type of sample. Mass tolerance for precursors in all data was±10 ppm, fragment mass tolerance for DDA data was 0.01 Da. Carbamidomethylation of cysteine residues was fixed modification, no variable modifications and no missed cleavages were allowed in MS1 search. For DDA data, the variable methionine oxidation and tmt10plex tags (where applicable) were allowed. The other search parameters were used by default.

**Table 4.** Summary on databases and target-decoy FDRs used to characterize different samples. *group-specific FDR, target-decoy approach.

| Sample type | NCBI:txid | Database | # proteins in fasta | FDR |
| --- | --- | --- | --- | --- |
| 19 single strains | all | SwissProt+TrEMBL |  | 0.05 |
| Fecal human microbiome | all | SwissProt+TrEMBL |  | 0.05* |
| Model I (*R. opacus* 1CP | 37919 | SwissProt+TrEMBL | 7769 | 0.05 |
| *R. erythropolis* X5 | - | Proteogenomic | 6407 |  |
| *G. alkanivorans* 135 | - | Proteogenomic | 5855 |  |
| *G. amicalis* 6-1 | 1220574 | SwissProt+TrEMBL | 4528 |  |
| *P. aryabhatai* 25) | 1358420 | SwissProt+TrEMBL | 6393 |  |
| Model II (*R. opacus* 1CP | 37919 | TrEMBL | 7769 | 0.05 |
| *R. opacus* 3D | - | Proteogenomic | 8870 |  |
| *R. opacus* S8 | - |  | 8102 |  |
| *R. erythropolis* X5 | - |  | 6407 |  |
| *R. qingshengii* 7B | - |  | 6277 |  |
| *R. qingshengii* VT6 | - |  | 6789 |  |
| *G. alkanivorans* 135 | - |  | 5855 |  |
| *G. polyisoprenivorans* 135) | - |  | 6248 |  |
| Model III (*R. opacus* 1CP | 37919 | SwissProt+TrEMBL | 7769 | 0.05 |
| *R. erythropolis* X5 | - | Proteogenomic | 6407 |  |
| *P. aryabhatai* 25 | - | Proteogenomic | 5855 |  |
| *G. amicalis* 6-1 | 1220574 | SwissProt+TrEMBL | 4528 |  |
| *G. alkanivorans* 135) | 1358420 | SwissProt+TrEMBL | 6393 |  |
| Single strain response 1<br>*R. opacus* 1CP | 37919 | SwissProt+TrEMBL | 7769 | 0.01 |
| Single strain response 2<br>*R. erythropolis* X5 | - | Proteogenomic | 6407 | 0.01 |

To estimate fold changes of microorganism biomasses between samples of model microbiomes I and II, protein identification was made against the pooled proteogenomic databases (**Table 4**). Quantitation was performed with DirectMS1Quant^40^.

To characterize changes in metabolic activity of *R. opacus* 1CP, the DirectMS1Quant was used for quantitation using the following parameters: differentially regulated proteins satisfy Benjamini Hochberg FDR < 0.05, fold change (FC) threshold was two standard deviations of log_2_FC distribution; intensity normalization by 1000 quantified peptides with maximal intensities was applied. Functional annotation and gene ontology analysis was performed using STRING db^41^. The annotation is publicly available at https://version-12-0.string-db.org/organism/STRG0A76PPK.

*R. erythropolis* X5 was quantified using Diffacto^42^ and QRePS^43^. Differentially regulated proteins were selected using the following criteria: |log_2_FC| > 1.2, |log_10_FDR| > Q3+1.5*IQR, where Q3 and IQR are 3rd quartile and interquartile range^43^. Functional annotation and gene ontology analysis were performed using STRING db. The annotation is freely available at https://version-12-0.string-db.org/organism/STRG0A44VNI. **Tables S1 and S2** containing quantitation results for *R. opacus* 1CP and *R. erythropolis* X5 are provided in supporting information.

## Results

### 1. Algorithm for two-stage blind database search identifies genus and species of a bacteria from LC-MS1 data

#### 1.1 Algorithm for two-stage search against SwissProt+TrEMBL database. The standard

DirectMS1 analysis using ms1searchpy search engine cannot handle protein databases with more than few hundred thousands of proteins. To solve that issue, we first used a preliminary search to identify the most probable strains presented in the sample using the whole bacterial database SwissProt+TrEMBL as input candidates. The algorithm for this search is based on a simple matching of theoretical peptide *m/z* values with experimental MS1 spectra. Based on the identifications from the preliminary search, the shortened database is composed of the most probable species. This shortened database is further used for accurate protein identification using the standard ms1searchpy search. Details on the algorithm are provided in the Supporting information (Appendix A, **Figure S1**). As an example, the runtime for full analysis on a computer with Intel(R) Core(TM) i7-7700HQ CPU @ 2.80GHz (4 cores) takes 4 hours: 3 hours for SwissProt+TrEMBL parsing and calculation of theoretical *m/z* (performed once), 6 minutes for preliminary search and 1 hour for standard search against the shortened database (a sum for 5 sample replicates).

#### 1.2 Two-stage blind search against SwissProt+TrEMBL demonstrates high accuracy in the identification of isolated bacterial species

To estimate the efficiency of the proposed two-stage identification approach, public LC-MS/MS data for 19 strain isolates was used^14^. The results of blind identification are presented in **Table 5**. The top identified microorganism matched the analyzed strain at the level of genus+species in 95 %, and 32 % of those cases matched the correct strain (“Level of match” column in **Table 5**). Analysis of top-3 identified microorganisms reveals that 11 of 19 cases showed the undoubted leader supported by at least 10 times higher numbers of identified proteins than the top-second and top-third taxons. The remaining eight cases demonstrated less than 2-fold difference in numbers of protein identifications distributed between the top-first and top-second organisms. This observation corresponds to the identification of taxons of the same species or species group and means a high similarity of proteomes. *E. coli* was the only strain identified at the level of the genus due to the classification within the ete3 toolkit^44^. The *E. coli* strain K-12 (NCBI:txid83333, 4518 proteins in SwissProt) is not reported with NCBITaxa().get_descendant_taxa() function for *E.coli* species (NCBI:txid562) which leads to it being excluded from the search space. At the next step, *E.coli* species database is also excluded in blind search due to its excessive size (1501705 proteins). Reportedly, differentiation between *Escherichia coli* and *Shigella*^45^ or *Bacillus cereus* and *Bacillus anthracis*^46^ is also problematic for MALDI TOF that requires development of specific solutions. The proposed blind search algorithm can be successfully applied for identification of individual species and strains using fast proteome profiling data.

**Table 5.** Summary of blind identification of bacteria against SwissProt+TrEMBL database shows correct match between the analyzed and top identified strains at the level of species groups, species and strains. Level of match was determined using phylogenetic trees available at https://phylot.biobyte.de/. Percentage is defined as (#proteins identified per species / total identified proteins)*100 %. Strains taken in multiple replicates were analyzed calculating the mean number of identifications across replicates.

| Strain analyzed | NCBI:txid | 1st taxon by #proteins, % | 2nd taxon by #proteins, % | 3rd taxon by #proteins, % | Level of match |
| --- | --- | --- | --- | --- | --- |
| *Acinetobacter baumannii*<br>DSM 30007 | 470<br>575584 | *Acinetobacter baumannii*<br>(45 %) | *Acinetobacter* sp.<br>FDAARGOS_559<br>(41 %) | *Acinetobacter* sp.<br>25977_8<br>(6 %) | strain |
| *Bacillus cereus*<br>DSM 31 | 1396<br>226900 | *Bacillus anthracis*<br>(51 %) | *Bacillus* sp. S66<br>(28 %) | *Bacillus* sp. B13<br>(5 %) | Species group |
| *Bacillus velezensis*<br>DSM 23117/FZB42 | 492670<br>326423 | *Bacillus velezensis*<br>FZB42 (32 %) | *Bacillus velezensis*<br>(19 %) | *Bacillus* sp.<br>VMFN-A1 (18 %) | strain |
| *Burkholderia cepacia*<br>ATCC 25416 | 292<br>983594 | *Burkholderia reimsis* (22 %) | *Burkholderia* sp.<br>LS-044 (16 %) | *Burkholderia lata*<br>(14 %) | Species group |
| *Burkholderia thailandensis*<br>DSM 13276 | 57975<br>271848 | *Burkholderia thailandensis*<br>(65-68 %) | *Burkholderia oklahomensis*<br>(6-8 %) | *Burkholderia* sp.<br>MSMB1552<br>(6-8 %) | strain |
| *Burkholderia thailandensis*<br>E125 | 57975 | *Burkholderia thailandensis*<br>(67-70 %) | *Burkholderia* sp.<br>MSMB1552<br>(8-12 %) | *Burkholderia oklahomensis*<br>(6-8 %) | species |
| *Burkholderia thailandensis*<br>E131 | 57975 | *Burkholderia thailandensis*<br>(67-68 %) | *Burkholderia* sp.<br>MSMB1552<br>(7 %) | *Burkholderia oklahomensis*<br>(1-9 %) | species |
| *Burkholderia thailandensis*<br>E153 | 57975 | *Burkholderia thailandensis*<br>(65-68 %) | *Burkholderia* sp.<br>MSMB1552<br>(7-13 %) | *Burkholderia oklahomensis*<br>(8 %) | species |
| *Burkholderia thailandensis*<br>LMG 20219 | 57975 | *Burkholderia thailandensis*<br>(67-71 %) | *Burkholderia* sp.<br>MSMB1552<br>(8-11 %) | *Burkholderia oklahomensis*<br>(1-8 %) | species |
| *Burkholderia oklahomensis*<br>DSM 21774 | 342113 | *Burkholderia oklahomensis*<br>(78-83 %) | *Burkholderia lata*<br>(4 %) | *Burkholderia* sp.<br>ABCPW 14<br>(3 %) | strain |
| *Citrobacter freundii*<br>DSM 30039 | 546<br>1006003 | uncultured<br>*Citrobacter* sp.<br>(57 %) | *Citrobacter freundii*<br>complex sp. CFNIH9<br>(14 %) | *Citrobacter* sp.<br>LUTT5<br>(8 %) | Species<br>group |
| *Enterococcus faecalis*<br>DSM 20371 | 1351 | *Enterococcus*<br>*faecalis*<br>(92 %) | *Enterococcus*<br>*faecium*<br>(2 %) | *Enterococcus*<br>*haemoperoxidus*<br>ATCC BAA-382<br>(1 %) | strain |
| *Mycobacteroides*<br>*abscessus*<br>DSM 44196 | 36809<br>561007 | *Mycobacteroides*<br>*abscessus* ATCC<br>19977 (57-58 %) | *Mycobacteroides*<br>*abscessus* subsp.<br>Abscessus (18-19 %) | *Mycobacteroides*<br>*franklinii*<br>(11 %) | strain |
| *Escherichia coli*<br>DSM 3871<br>(K12 W3110<br>derivative) | 562<br>83333 | *Escherichia*<br>*fergusonii* ATCC<br>35469 (20 %) | *Escherichia* sp.<br>KTE114<br>(18 %) | *Shigella*<br>*boydii*<br>(12 %) | genus |
| *Pseudomonas*<br>*aeruginosa*<br>ATCC 27853 | 287 | *Pseudomonas*<br>*aeruginosa*<br>(93-94 %) | *Pseudomonas* sp.<br>ATCC 13867<br>(2 %) | *Pseudomonas*<br>*nitroreducens*<br>(1 %) | species |
| *Staphylococcus*<br>*epidermidis*<br>DSM 1798 | 1282 | *Staphylococcus*<br>*epidermidis*<br>(85 %) | *Staphylococcus* sp.<br>HMSC34G04<br>(4 %) | *Staphylococcus* sp.<br>RIT622<br>(2 %) | species |
| *Staphylococcus*<br>*aureus* DSM<br>20231/NCTC 8532 | 1280<br>1241616 | *Staphylococcus*<br>*aureus*<br>(95 %) | *Staphylococcus*<br>*pasteuri*<br>(1 %) | *Staphylococcus*<br>*argenteus*<br>(1 %) | species |
| *Vibrio cholerae*<br>NIH 41 | 666 | *Vibrio cholerae*<br>(77 %) | *Vibrio mimicus*<br>(8 %) | *Vibrio metoecus*<br>(6 %) | species |
| *Yersinia*<br>*pseudotuberculosis*<br>DSM 8992 | 633 | *Yersinia*<br>*wautersia*<br>(41 %) | *Yersinia*<br>*pseudotuberculosis*<br>(28 %) | *Yersinia*<br>*similis*<br>(5 %) | species |

### 2. Two-stage blind database search identifies relative composition of microbial community at level of genus

#### 2.1 Model mixtures of soil bacteria

To study the performance of a two-stage blind search to identify microbiome composition, the soil bacteria mixed at known concentrations (**Tables 1–3**) were used. The main motivation was to investigate how blind searches resolve the metaproteomic samples containing different species of the same genus. The results of blind identification of sample compositions are summarized in **Table 6**. The actual and estimated compositions of each model microbiome were compared at two levels: genus and species. In blind identification, the mean difference between the actual and identified components of microbiome composition was 12 % at the level of genus. The difference between the actual and estimated compositions depended on the microorganisms under study. In our example, the content of *Rhodococcus* was easily identified and often overestimated, while *Gordonia* was identified in all samples (even though its content was below 15 %), but always underestimated. *Priestia* was identified with high accuracy within a few percent, but only if its actual fraction in the sample was higher than 30 %. Below 30 %, *Priestia* was not identified at all. At the level of species, the mean difference between the estimated and actual content was 15 %. However, the number of missing identifications of microorganisms increased from 20 % at the level of genus to 40 % at the level of species. Thus, the current realization of blind search algorithm for identification of microbiome composition provides the most complete information at the level of genus.

**Table 6.**
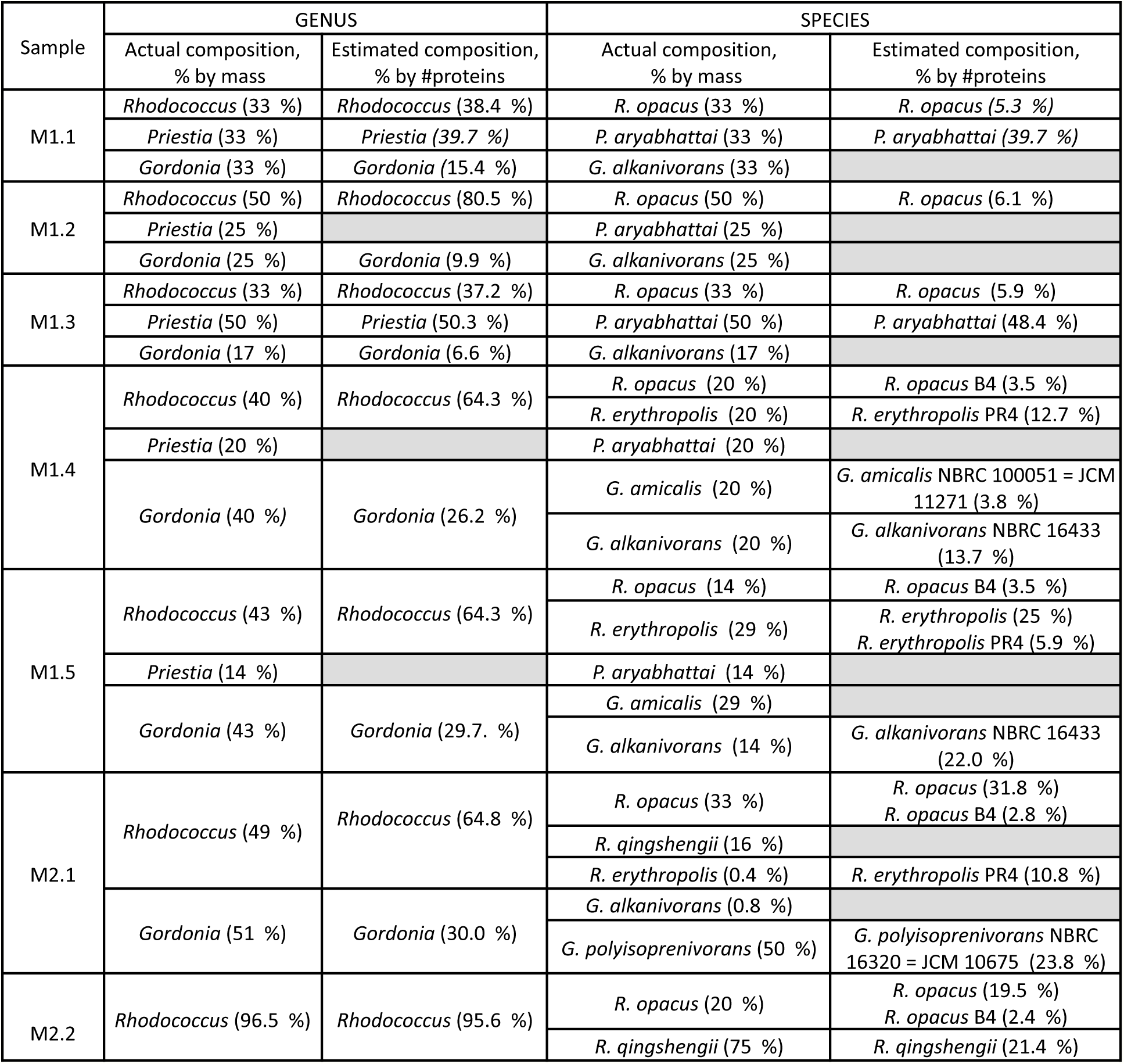
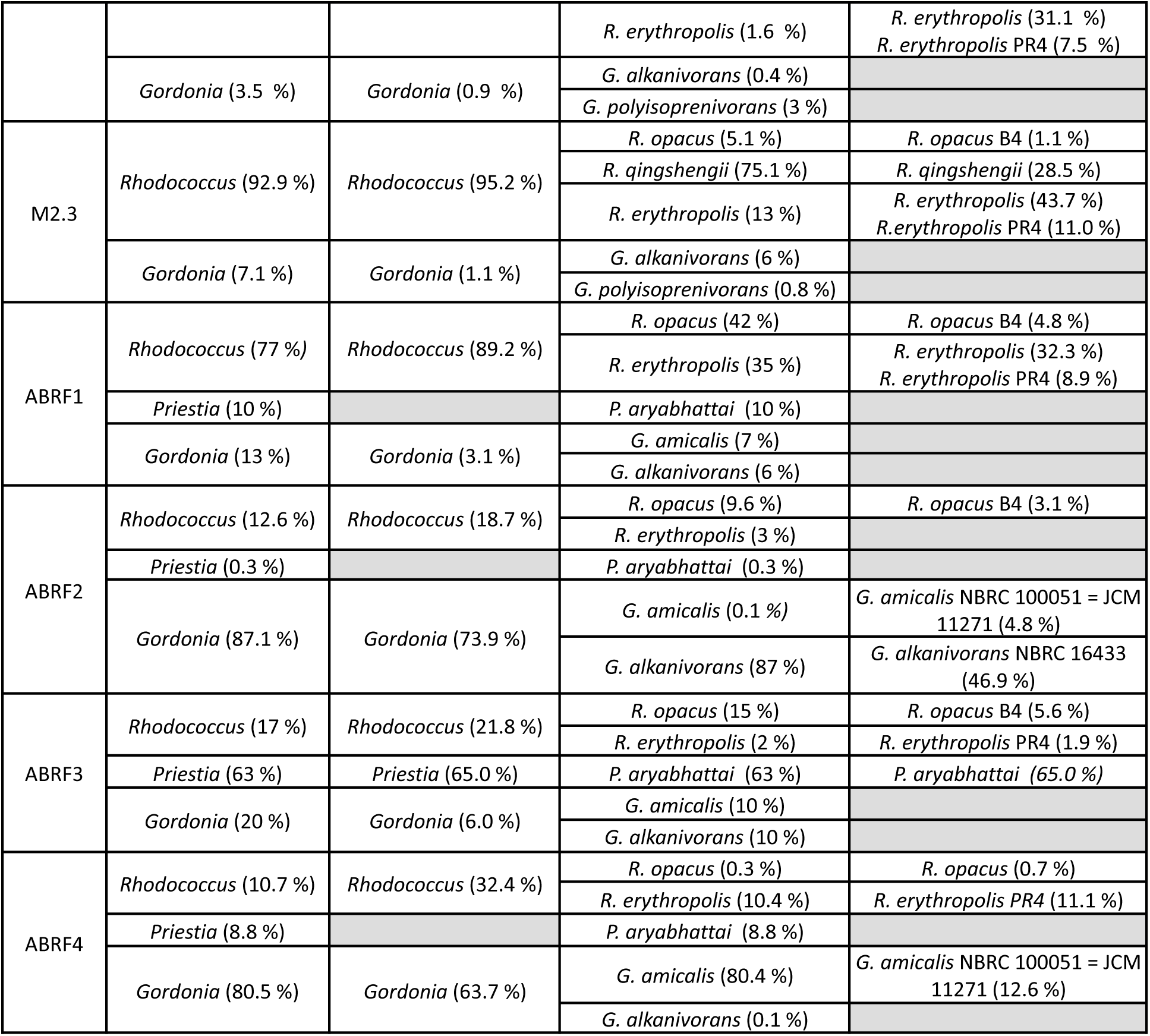
Summary of blind identification of microbial compositions against SwissProt+TrEMBL database. Sample labels and actual compositions correspond to Tables 1–3. Percentage in actual compositions is defined as (peptide mass per genus (species) / total peptide mass)*100 %. Percentage in estimated compositions is defined as (#proteins per genus (species) / total #proteins)*100 %. Empty cells mean missing identification of microorganisms from a given genus or species. The mean difference between actual and estimated composition was calculated as ∑|Actual % - Estimated %|/N, where N=33 is the number of rows in the column “actual composition, % by mass”. Missings were imputed by zero.

#### 2.2 Fecal microbiome

To test the algorithm in identifying the composition of the real microbiome, the public dataset for human fecal microbiome was used^14^. In the referenced publication, the sample was analyzed using a multi-omics approach involving metagenomics, metatranscriptomics, and metaproteomics. Taxonomic profiling of the results obtained using these methods has shown that the families *Ruminococcaceae*, *Lachnospiraceae, Eubacteriaceae* and *Clostridiaceae* were the most abundant. Among the organisms identified using the blind search (**Figure 1**), the following bacteria were found: (1) *Ruminococcaceae* family: *Faecalibacterium prausnitzii*, *Ruminococcus* sp. AM54-14NS, *Faecalibacterium* sp. AF28-13AC, *Ruminococcus* sp. AM29-26, *Ruminococcus* sp. AF31-14BH; (2) *Eubacteriaceae* family: *Eubacterium rectale* CAG:36; (3) *Lachnospiraceae* family: *E. rectale* ATCC 33656; and (4) *Clostridiaceae* family*: Clostridia* sp. AF34-10BH. Despite the low number of identified proteins for the sample, it was possible to identify four families composing approximately 90 % of the sample (according to the data from^14^) and suggest a genus composition for them. Thus, the proposed algorithm can be applicable for the identification of both individual strains and complex samples such as the real human gut microbiome.

**Figure 1.**
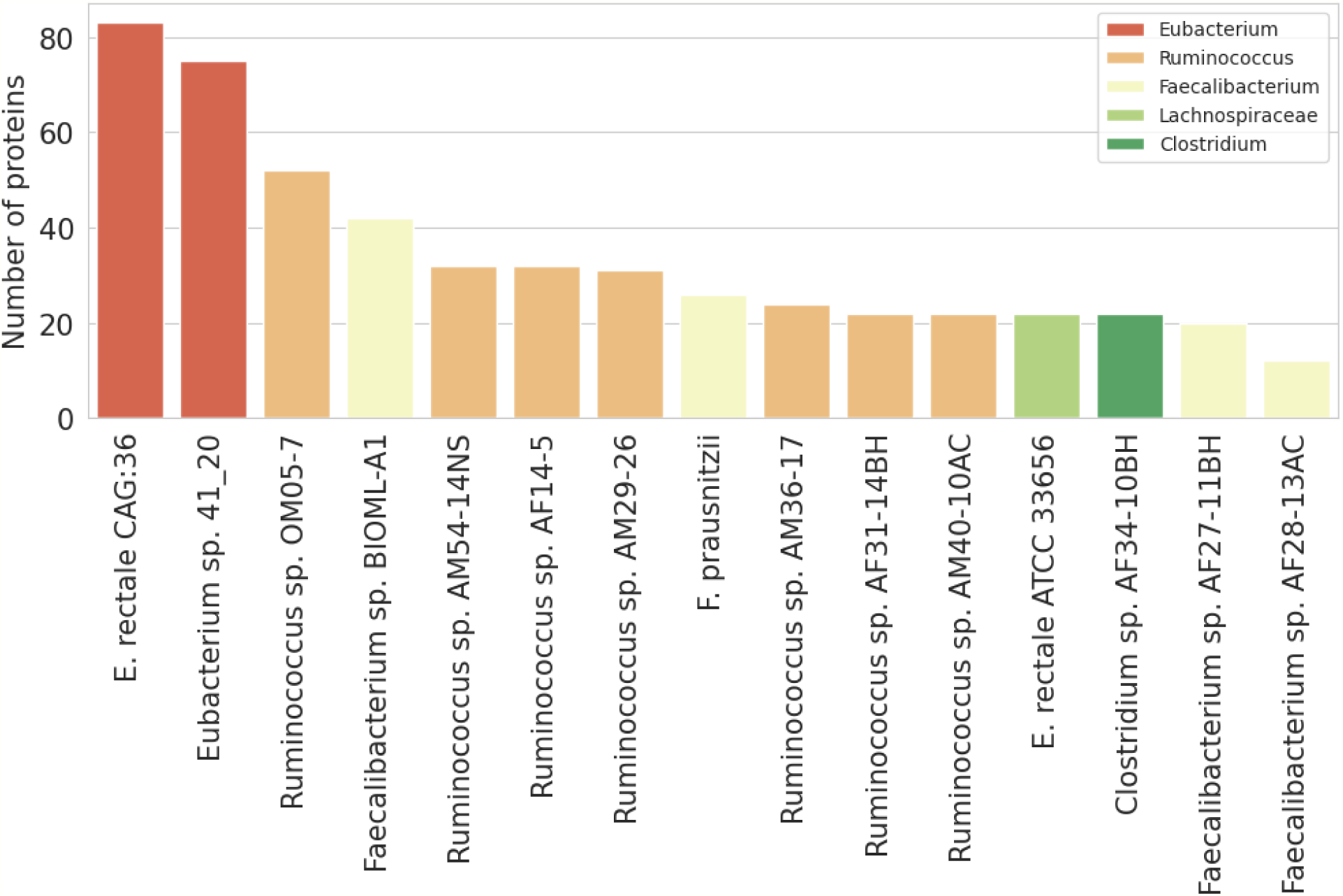
Blind identification of fecal microbiome composition using DirectMS1. Organism-specific FDR is 0.05.

### 3. DirectMS1 allows correct determination of the fold change of species biomass between microbiome samples

Model microbiomes I, II and III (**Tables 1-3**) were used to analyze strain abundance variation between different samples with the following procedure. After quantitation at peptide level, the fold changes of all quantified peptides without decoys were used to plot the baseline distribution (**Figure 2a**). Then, fold change distribution was built for all peptides from each strain of the model microbiome (“strain” distribution). The baseline was subtracted from the “strain” distribution and the corrected distribution was analyzed (**Figures 2 b,c,d**). A weighted average of fold changes for the corrected distribution provides an approximation of the biomass fold change for the selected strain, where weights are non-negative normalized densities.

**Figure 2.**
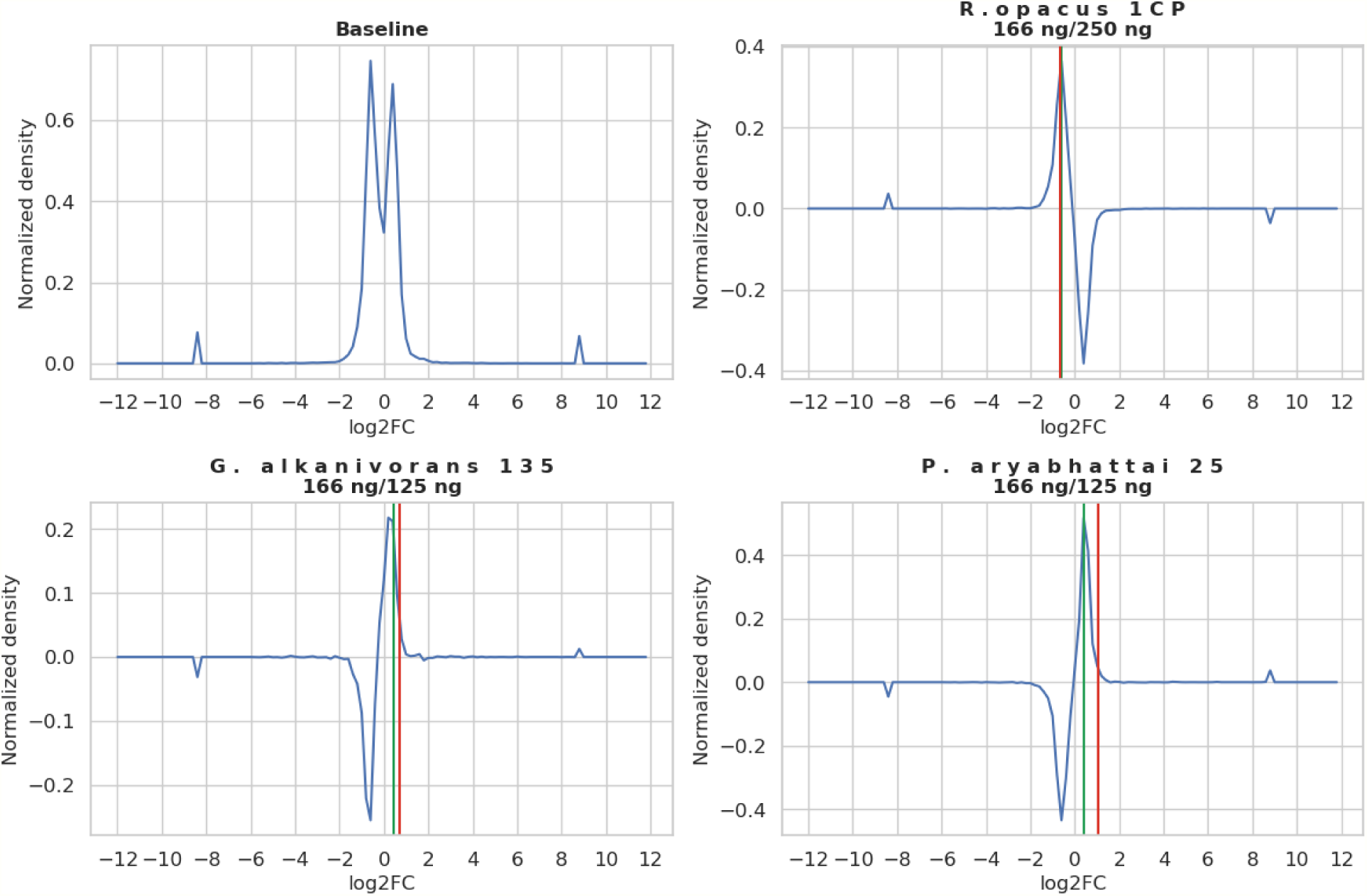
The normalized densities of log_2_FC distribution for M1.1/M1.2 comparison. The green line stands for the actual ratio of peptide masses (given in subtitles) and the red line stands for the calculated ratio, log2-scaled. FC is the fold change in strain biomass between model microbiomes.

**Figure 3** summarizes estimates of fold changes in strain biomass between samples and compares it to actual mass ratios. For all samples from the Model I, the experimentally measured fold changes coincide well with the actual values (Pearson correlation coefficient of 0.97, **Figure 3a**). It means that species mixed at ratios 1:1, 1:2, 1:3 and composing at least from 15 to 20 % of a microbiome are typically well resolved with our method. For Model II and III, a decrease in fold change predictions was observed (**Figure 3b,c**). Model II represented the extreme case of uniform composition of the same species (*R. opacus* 1CP, *R. opacus* 3D, *R. opacus* S8; *R. erythropolis* X5, *R. qingshengii* 7B, *R. qingshengii* VT6, *G. alkanivorans* 135, *G. polyisoprenivorans* 135) mixed in a range of concentrations differing by up to two orders (**Table 2** in M&M). This complex case resulted in a decrease of Pearson correlation to 0.81 due to *Gordonia* species composing just 3.5 % (M2.2) and 7.1 % (M2.3) of the model microbiome (**Figure 3b**).

**Figure 3.**
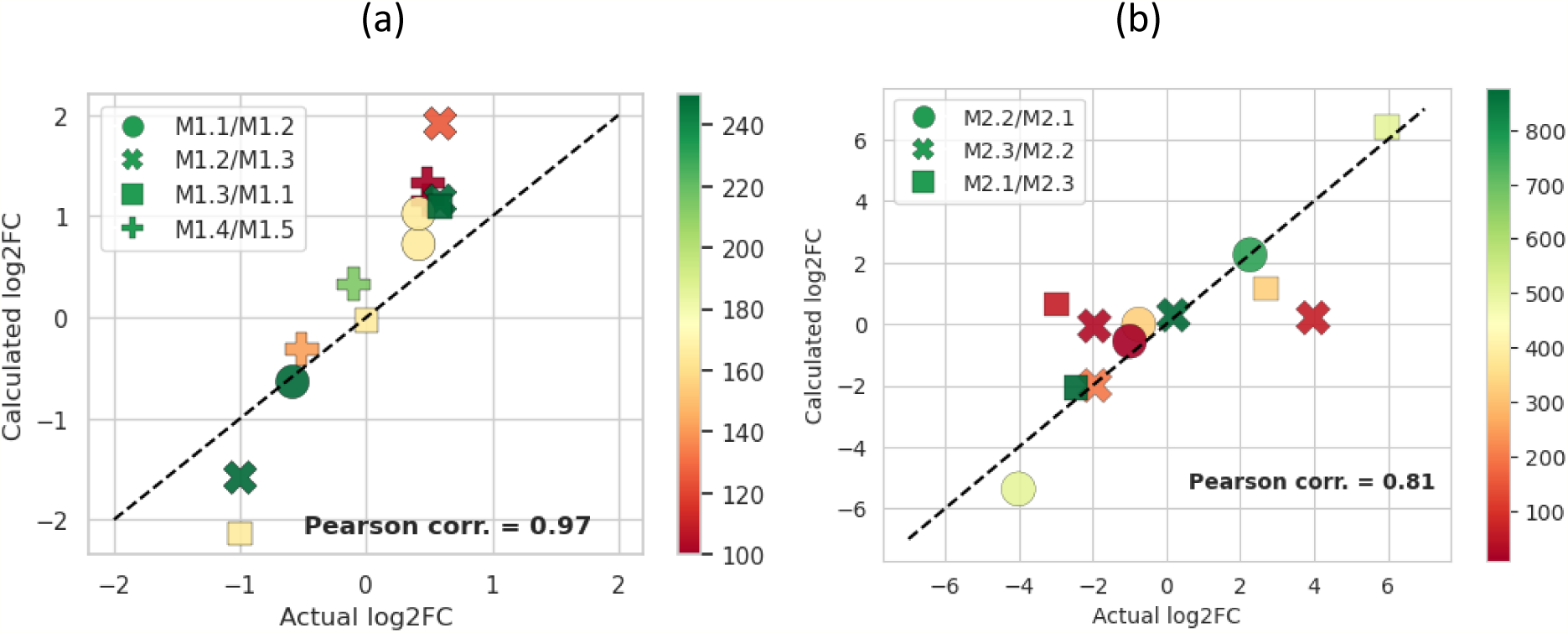
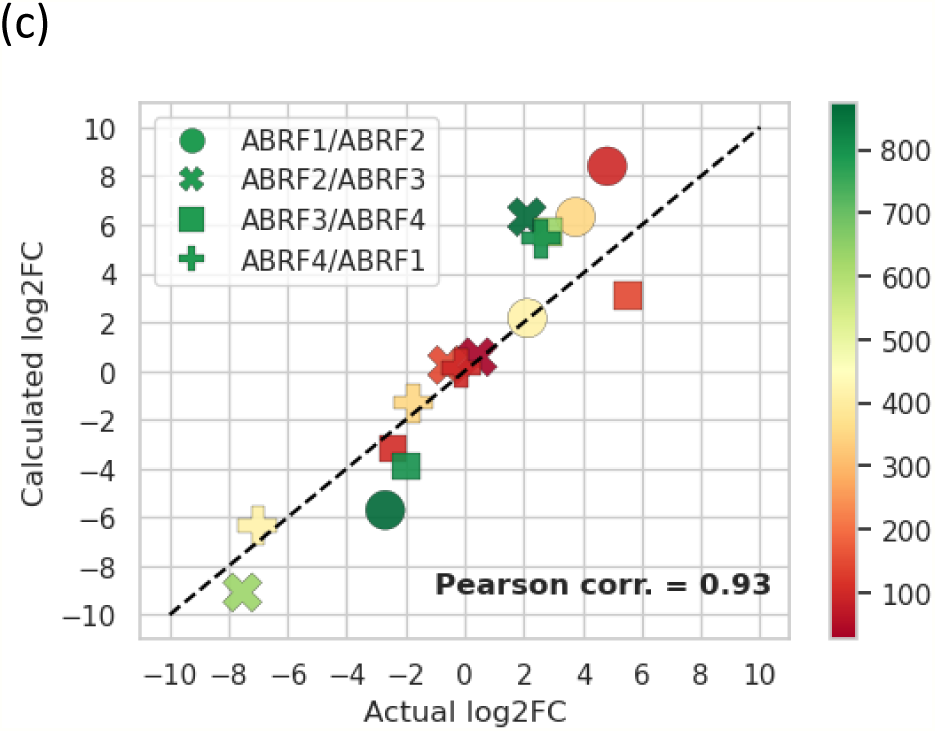
Correlation of actual and experimentally measured fold changes in strain biomass for Model I (a), Model II (b), Model III (c). Y: fold change estimate from experimental data, X: actual ratios, in log2-scale. Legends correspond to labels in Tables 3–5. The color corresponds to the maximum of two compared species biomasses, in nanograms.

Model III had compositions differing at the level of species (*R. opacus* 1CP, *R. erythropolis* X5, *P. aryabhattai* 25, *G. amicalis* 6-1, *G. alkanivorans* 135) (**Table 3** in M&M) and covering the similar range of concentrations as samples from Model II. Model III is less complex compared with Model II that results in more accurate fold change estimations (R = 0.93, **Figure 3c**).

For the low amounts (species or genera content within a few percent) in one of the samples, the missing values can occur and subsequent imputation of peptide abundance results in a decreased accuracy. **Figure S2** illustrates such a case with the bacterium *P. aryabhattai* 25 from Model III ABRF2/ABRF3, for which the peptide content was 3 ng and 631 ng, respectively. If the total biomass of microorganisms is low in both samples, the method could not quantify enough peptides to construct the distribution. This case is illustrated with the bacterium *R. erythropolis* X5 in **Figure S2**, for which there was no fold change estimation provided.

Another case we explored is the fold change predictions for mixtures with high overlap of the identified peptides coming from different species. For example, the fold change distributions for *G. alkanivorans* 135 and *G*. *amicalis* 6-1 from Model III ABRF2/ABRF3 coincide (**Figure S2**) and the fold change estimate matches the sum of the peptide masses of both strains. This trend was observed for all mixtures analyzed. Next example is the Model II M2.2/M2.3 comparison demonstrating an absence (or undetectable concentrations) of unique peptides required for distinguishing the *R. opacus* strains, as well as *R. erythropolis* and *R. qingshengii* species (**Figure S3**). From a practical viewpoint, it means that the measured fold change between the components of complex microbiomes matches the sum of the biomasses of all strains identified by the shared peptides.

### 4. DirectMS1 allows the characterization of changes in metabolic pathways and biodegradation ability of strain isolates

The performance of DirectMS1 to characterize metabolic response of bacteria to the stimulus was tested using two pollutant-degrading actinobacteria: (1) *Rhodococcus opacus* 1CP grown in presence of glucose, benzoate, phenol, and 4-chlorophenol; and (2) *Rhodococcus erythropolis* X5 grown at 28 °C and 6 °C in presence of *n*-hexadecane. **Figure 4–5** summarizes the results of quantitation analysis.

**Figure 4.**
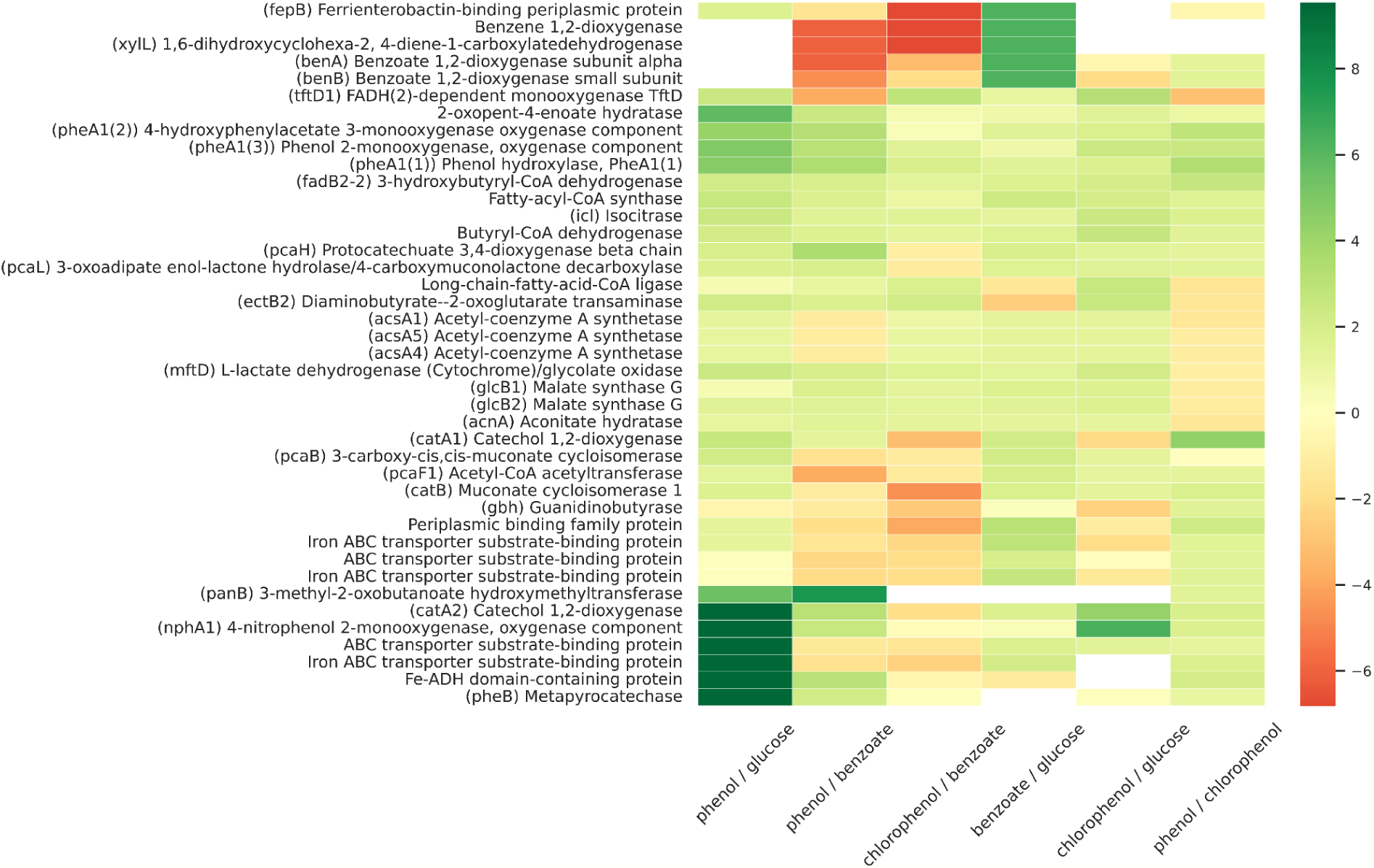
Proteomic response of actinobacteria *Rhodococcus opacus* 1CP to the presence of aromatic compounds: benzoate, phenol, and 4-chlorophenol, compared with glucose. Quantitation: DirectMS1Quant^40^ [<u>10.1021/acs.analchem.2c02255</u>]. Protein selection corresponds to the most enriched biological processes (GO score ≥ 6).

**Figure 5.**
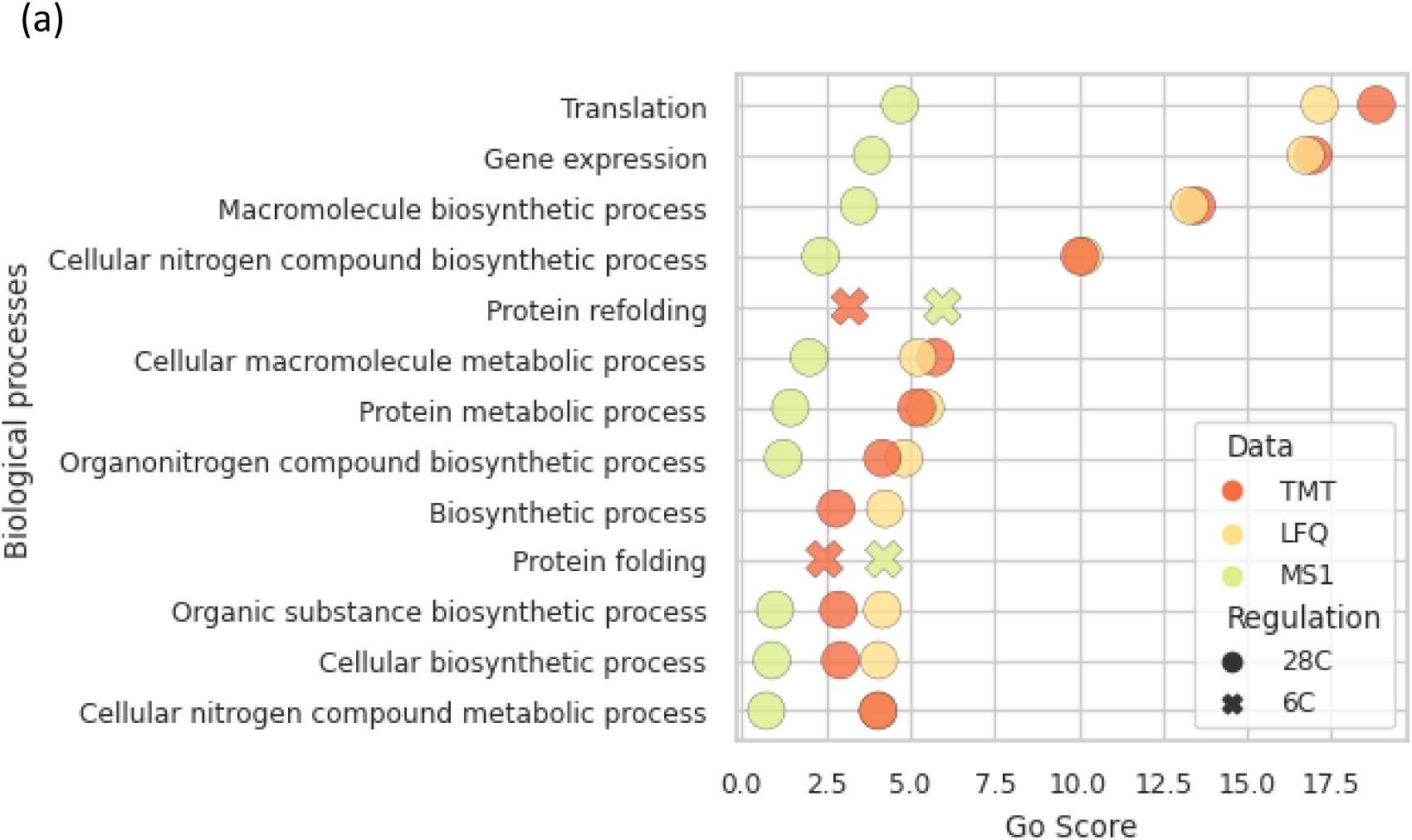
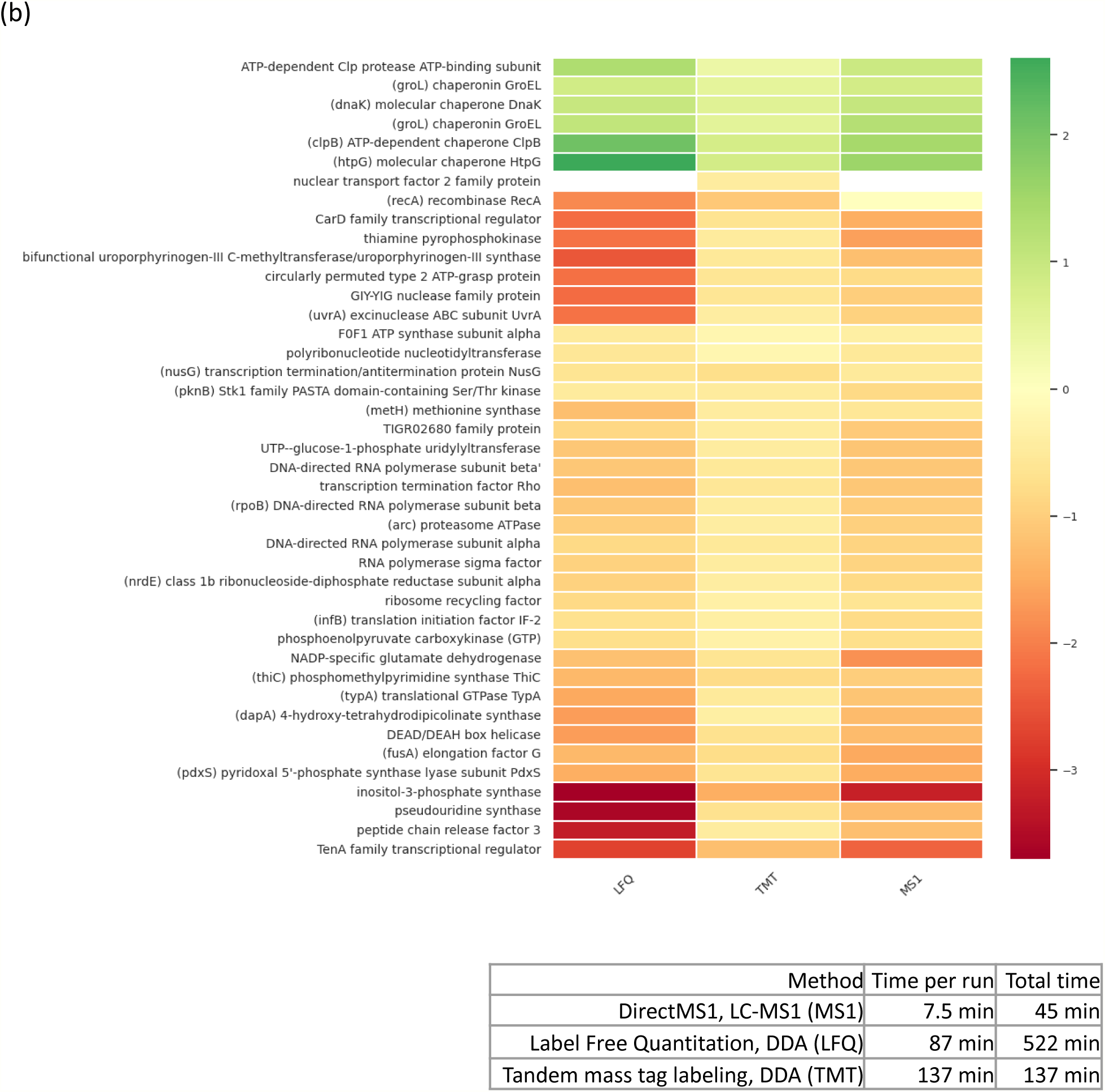
Response of actinobacteria *Rhodococcus erythropolis* X5 grown on *n*-hexadecane to temperature change from 6 °C to 28 °C. Quantitation: Diffacto^42^ (LFQ, MS1-only acquisition), MSFragger+Scavager+NSAF (LFQ, DDA), and MSFragger+Scavager+Diffacto (DDA, TMT labeling). GO Score = E * log10 fdr, where E is the enrichment of biological processes, fdr is the statistical significance of the GO enrichment corrected for multiple comparisons using Benjamini-Hochberg method^52^.

**Figure 4** summarizes the key differences in the proteomes of *Rhodococcus opacus* 1CP grown in the presence of phenol, 4-chlorophenol and benzoate, compared with glucose. *R. opacus* 1CP is the actinobacterium able to degrade phenol and their derivatives^34^. In our study, the presence of phenol resulted in upregulation of enzymes involved in phenol degradation: three different phenol hydroxylases ((**pheA1(1**), **pheA1(2**), and **pheA1(3**)), and 2-oxopent-4-enoate hydratase (**R1CP_00930**). These enzymes are also induced in the presence of benzoate and 4-chlorophenol, but with smaller magnitude. The data obtained are in full agreement with previously obtained real-time PCR results^47^. Using specific primers for the small subunit of all three phenol hydroxylases of strain 1 CP growing in the presence of phenol, the gene activation has been shown; an increase for phe A1(3) was approximately 2000 times.

Enzyme cluster consisting of upregulated 3-methyl-2-oxobutanoate hydroxymethyltransferase (**panB)**, catechol 1,2-dioxygenases (**catA1/catA2)**, 4-nitrophenol 2-monooxygenase (**nphA1)**, metapyrocatechase (**pheB)**, and Fe-ADH domain-containing protein were specific to phenol. For degradation of 4-chlorophenol, we observed no specific features, the upregulated enzymes were the same as for phenol conditions. For benzoate conditions, we found specific pattern of enzymes degrading benzoate: benzene 1,2-dioxygenase (**A8787_2660)**, benzoate 1,2-dioxygenase subunit alpha (**benA)**, benzoate 1,2-dioxygenase small subunit (**benB)**, 1,6-dihydroxycyclohexa-2, 4-diene-1-carboxylatedehydrogenase (**xylL)**, and ferrienterobactin-binding periplasmic protein (**fepB)**.

In total, the results of ultrafast proteomics profiling were consistent with the expected changes in metabolic activity and can serve as a valuable support in characterization of pollutant degradation activity of different microorganisms.

**Figure 5** shows response of the bacterium *Rhodococcus erythropolis* X5 grown on *n*-hexadecane to temperature change from 6 °C to 28 °C. *R. erythropolis* X5 is the psychrotrophic bacterium that possesses a wide range of catabolic activities within 4°C to 28°C temperature range. Specifically, this strain can degrade *n*-alkanes which constitute the main fraction of oil pollution^25^. In our study, the proteome of X5 strain grown on *n*-hexadecane at 6 °C was compared with its proteome at 28 °C. At 6 °C, we observed proteins related to all steps of *n*-alkane degradation including oxidation to alcohols (QEX08599, QEX09356, etc), oxidation to aldehydes (QEX08400, QEX09399, etc), oxidation to fatty acids and fatty acid metabolism (QEX08594, QEX12285, QEX08597, etc), exopolysaccharide production (QEX09055, QEX10519, QEX12651, QEX11399, etc), and iron transport (QEX13740, QEX13741, QEX09379, etc) (**Table S2, Appendix B**). Besides enzymes involved in *n*-alkane biodegradation, the chaperons/chaperonins (QEX09859, QEX13151, QEX09986, QEX09823) and stress/protection proteins (QEX09757, QEX11651, QEX11169) were upregulated at 6 °C. Also, a significant loss of 30S/50S ribosomal proteins was observed at cold conditions. Numerous studies report that loss of ribosomal proteins in bacteria is the reaction to stress and is necessary for survival under nutrient- or energy-limited conditions^48,49,50,51^. GO analysis revealed that translation, gene expression, and protein folding were significantly affected at cold conditions (**Figure 5a**). Proteins involved in DNA and RNA processing, regulation of transcription and translation are shown in **Figure 5b** and **Appendix B.**

At 28 °C, we observed the abundant protein group corresponding to degradation of *n*-alkanes, but expressed from other parts of genome: *n*-alkanes to alcohols (QEX08601, QEX08447), alcohols to aldehydes (QEX10266, QEX10225, QEX10036), aldehydes to fatty acids / fatty acid metabolism (QEX12404, QEX10265, QEX10256, and many others), exopolysaccharide production (QEX12160, QEX13966, QEX14217, etc), and iron transport (QEX11646, QEX12370, QEX11669, etc) (**Table S2, Appendix B**). This observation suggests that different parts of the *R. erythropolis* X5 genome were transcribed to translate the enzymes needed for *n*-alkane degradation under optimal and cold conditions. This transcriptional behavior can be a conserved mechanism of coping with stress.

To confirm the technical reproducibility of the results, a comparison was made between label-free DirectMS1, DDA LFQ and TMT-based quantitation. **Figure 5a** shows the enriched biological processes obtained using the three quantitation methods. The results of the analysis show that the set of enriched biological processes for all methods has a high overlap. The protein content behind the top GO enrichments, measured using different quantitation workflows, is shown in **Figure 5b**. Legend in **Figure 5b** compares the duration of the experimental methods for the used quantitation methods. DirectMS1 requires less experimental time providing comparable performance.

## Discussion

We demonstrate that the DirectMS1 method for fast proteomic profiling can be effectively used for functional characterization of strain isolates and quantitative assessment of microbial communities. Our algorithm for two-stage blind database search discriminated bacteria to species level from LC-MS1 data with high accuracy of 95 %, and 32 % of those cases were matched to the correct strains. The algorithm identifies the composition of microbial communities at the level of genus and provides quantitative estimates in strain biomass. DirectMS1 for metaproteomics takes a position between MALDI TOF MS and LC-MS/MS-based proteomics, combining both fast identification and quantitative assessment of strain isolates and microbiomes at reasonable costs.

Besides MALDI-TOF MS, 16S rRNA gene sequencing and whole-genome sequencing are widely used for microorganism identification^53^. The accuracy of microorganism identification depends on the number of strain-specific features measured: the larger the number of features, the higher the level of discrimination. For example, 16S rRNA gene sequencing can possess limits in discriminating to the species level^54,55^. MALDI-TOF MS is often positioned as a more accurate technique capable of demonstrating high discrimination to the species and even strain levels^56^. For example, *Citrobacter* strains are identified to the species level by the MALDI-TOF MS, while 16S rRNA gene sequencing could not discriminate them clearly^54^. Another example, *Yersinia pestis* and *Yersinia pseudotuberculosis* are difficult to separate^57^. These species have similar genomes and identical 16S rRNA genes^58^, and their differentiation by MALDI-TOF MS is also challenging^46^. *Yersinia pseudotuberculosis* and *Yersinia wautersii* are also genetically close^59^. *Bacillus cereus* and *Bacillus anthracis*^46^, *Escherichia coli, Escherichia fergusonii* and *Shigella*^60,61,45^ are also represent challenges in discrimination. These earlier findings agree well with the top-ranking identifications reported during the blind search of strain isolates (**Table 5**). From the literature, we see that some species largely sharing genomes can generate identical 16S rRNA and very similar proteome profiles, and then can be resolved by only a combination of methods including whole genome sequencing.

The 5-minute LC-MS1-based proteome profiling of strain isolates responding to environmental conditions demonstrated high performance in quantitation (**Figures 4-5**). We detected the key enzymes degrading aromatic compounds, such as phenol, benzoate and 4-chlorophenol, that are in full agreement with previous studies^62,63^. We measured the key enzymes involved in the oxidation of *n*-alkanes to alcohols (i.e. alkane monooxygenases and cytochrome P450 family proteins); alcohols oxidation to aldehydes (alcohol dehydrogenases); aldehydes oxidation to fatty acids (aldehyde dehydrogenases); fatty acid metabolism (acyl-CoA dehydrogenases, fatty acid-CoA synthases, fatty acid synthesis proteins); exopolysaccharide production (glycosyltransferases, glucose dehydrogenases, fructose synthases, mannose dehydrogenases, etc.); and iron transport (ferredoxins, ferredoxin reductases, ABC transporter proteins)^64^. We conclude that DirectMS1 was proved as a valuable approach to functional annotation of strain isolates responding to the environment.

## Acknowledgements

I.T. and M.G. thank Prof. Victor Zgoda from “Human Proteome” Core Facility at the Institute of Biomedical Chemistry for help with implementation of DirectMS1 method on the Orbitrap FTMS system.

## Data availability statement

LC-MS1 and LC-MS/MS data are available at proteomexchange.org (PXD050587, PXD050761, PXD050807, PXD050887). Code for blind search, identification of microbial sample composition and assessment of species biomass change between samples is available at GitHub (https://github.com/kazakova/Metaproteomics-DirectMS1).

## Supporting information

Supporting information is available (Appendix A-B, Figures S1, S2, S3, Tables S1 and S2).

## Funding

The study (developing DirectMS1 and bioinformatic solutions for metaproteome profiling, study design, sample preparation, data processing, etc) was supported by the Russian Science Foundation grant No. 20-14-00229.

F.K. and V.G. acknowledge generous grants to the VILLUM Center for Bioanalytical Sciences (VILLUM Foundation grant no. 7292), PRO-MS: Danish National Mass Spectrometry Platform for Functional Proteomics (grant no. 5072-00007B), and the Novo Nordisk Foundation (INTEGRA, NNF20OC0061575) for support of instrumentation infrastructure at the University of Southern Denmark.

## Author contributions

Conceptualization, I.T.; Methodology, E.K.,J.B., M.I., Y.D., I.S., and I.T.; Software, M.I. and E.K.; Validation, E.K., J.B., and Z.Z.; Formal Analysis, M.I. and E.K.; Investigation, E.K., T.K., J.B., V.P., K.P, V.G., F.K., Y.D., I.S.; Data Curation, T.K. and E.K.; Resources, E.K., T.K., J.B., Y.D., I.S.; Writing – Original Draft Preparation, I.T.; Writing – Review & Editing, I.A., M.I., V.G., F.K, M.G.; Visualization, E.K.; Supervision, I.T.; Project Administration, I.T., M.G.

## Notes

### Competing Interest Statement

The authors have declared no competing interest.

### Summary of Updates

Changes in Title, Abstract and in the manuscript text

