## Supporting Information for "Ultrafast metaproteomics for quantitative assessment of strain isolates and microbiomes"

**Content**

**Appendix A.** Description of the algorithm behind two-stage database search for blind identification of bacteria using LC-MS1 data.
**Figure S1.** Scheme for algorithm behind two-stage database search for blind identification of bacteria using LC-MS1 data.
**Figure S2.** The normalized densities of $log_2$FC distribution for ABRF2/ABRF3 mock microbiome comparison. The green line stands for theoretical value, calculated from peptide masses (shown as plot subtitles).
**Figure S3.** The normalized densities of $log_2$FC distribution for M2.2/M2.1 comparison. The green line stands for theoretical value, calculated from peptide masses (shown as plots subtitles).
**Appendix B.** Protein signatures involved in degradation of n-alkanes at 6 $^0$C and at 28 $^0$C.

**Tables S1 and S2** contain quantification results for *R.opacus* 1CP and *R. erythropolis* X5 and are provided as the separate .xlsx files.

**Appendix A.** *Description of the algorithm behind two-stage database search for blind identification of bacteria using LC-MS1 data.*

Our first step was to develop an algorithm for MS1-based identification of the bacterial species when no prior knowledge of the bacteria in the sample is available. Such an algorithm requires the peptide-spectrum matching against the whole bacterial database SwissProt+TrEMBL whose current size is 62 Gb and contains more than 160 million of protein sequences. To keep execution time feasible, multiple optimization steps were applied.

First, the UniProt database was divided into 104758 organism-specific databases (101656 TrEMBL+SwissProt and 3102 SwissProt) using taxonomic identifiers (https://www.uniprot.org/help/taxonomic_identifier) from protein descriptions and number of proteins in each database was calculated (Fig S1-I). For each taxonomic identifier the rank was determined with NCBITaxa().get_rank() function from ete3. In the following analysis only organisms with strain, subspecies, forma specialis, isolate, serotype or serogroup rank were considered (99986 identifiers in total). Then with NCBITaxa().get_descendant_taxa() and NCBITaxa().get_lineage() each identifier was assigned to the group containing the species identifier and all the related to this species identifiers with lower ranks (85486 groups in total) (Fig. S1-II). The next step was to determine the group leader for each group i.e. the database containing the highest number of proteins (Fig. S1-IIIa). If some of the organisms from the group were present in the SwissProt database, group-leaders were obtained independently from both SwissProt and TrEMBL+SwissProt (Fig. S1-IIIb). This part was crucial in the cases when TrEMBL+SwissProt group leader was a joint database containing dozens of thousands of proteins and therefore will unlikely provide any reasonable final results due to the extremely large search space. As a result of this optimization the number of organism-specific databases was reduced from 101656 to 86058. The leaders whose name contains ‘sp.’ or ‘uncultured’ were excluded for analysis of mock microbiomes which leads to 27929 protein databases in the remnant (Fig. S1-IV).

After the group leaders’ selection a reduced protein database was prepared for the preliminary analysis. First, all species with less than 200 proteins were excluded from the analysis. Second, all SwissProt proteins and 2000 randomly chosen TrEMBL proteins were added to the reduced database for each bacterial species. After all, the reduced database size was 19 Gb which means ~70% reduction in the total number of proteins.

At the next step, the reduced protein database was theoretically digested into tryptic peptides with 0 missed cleavages and a length range of 9 to 15 amino acid residues. Neutral masses were calculated for every peptide and were stored as unique sets of values per protein (Fig. S1-V). Next, calculated masses were matched against peptide isotopic cluster features detected using biosaur2 software. Only the peptide features with 2+ or 3+ charges and minimal three visible carbon isotopes were used. Matching was done using an initial 4 ppm mass accuracy. Those strict limitations in peptide length, charges and isotope number were applied only in this preliminary step to reduce computational resources and execution time and the following DirectMS1 analysis was done using standard parameters described in the manuscript. Matching scores were calculated for every protein using a binomial model similar to the protein scoring concept used in DirectMS1 approach. The score was defined using equation:

$$score_i = -\log_{10}( SF (k_i-1, n_i, p) ) \quad (1),$$

where SF is survival function for binomial distribution, $k_i$ is the number of matched neutral masses for protein i, $n_i$ is the total number of neutral masses for protein i and p is the probability of random mass match.

p was calculated as the ratio between the number of matched neutral masses to the total number of neutral masses used for search.

At the next step, the proteins with scores less than 4 (0.01% probability) were removed. Differences between theoretical and experimental masses for species with the highest number of matched proteins were fitted using sum of normal and uniform distributions to extract mean shift and standard deviation of normal distribution. Mean +- 3 standard deviations was used as new final mass accuracy and the search was done again (Fig. S1-VI). Finally, 15 species with maximal number of proteins with scores above 4 were chosen and used in the standard DirectMS1 analysis using ms1searchpy search engine using all available SwissProt and UniProt proteins (Fig. S1-VII).

**Figure S1.** Scheme for algorithm behind two-stage database search for blind identification of bacteria using LC-MS1 data.

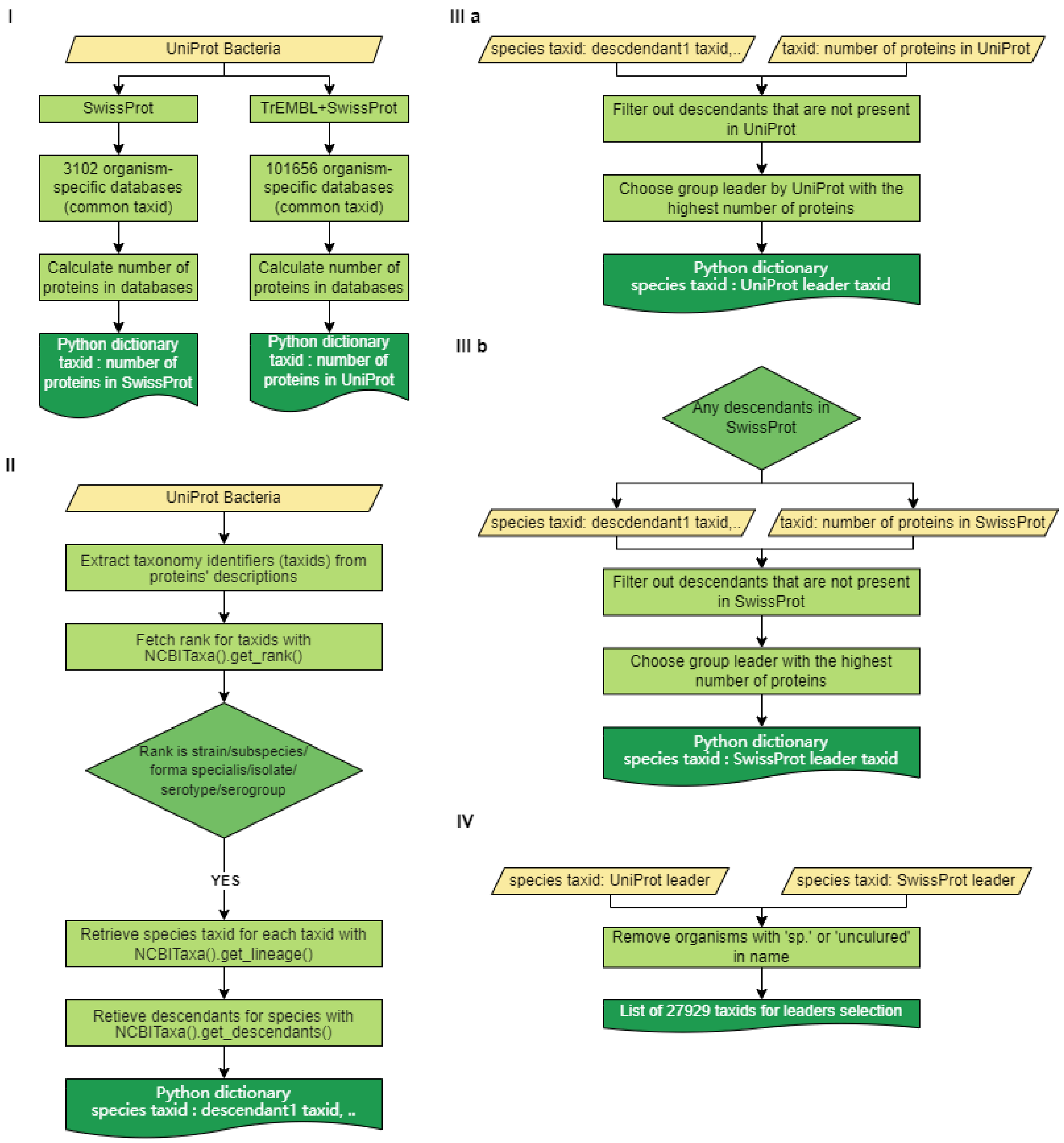

V

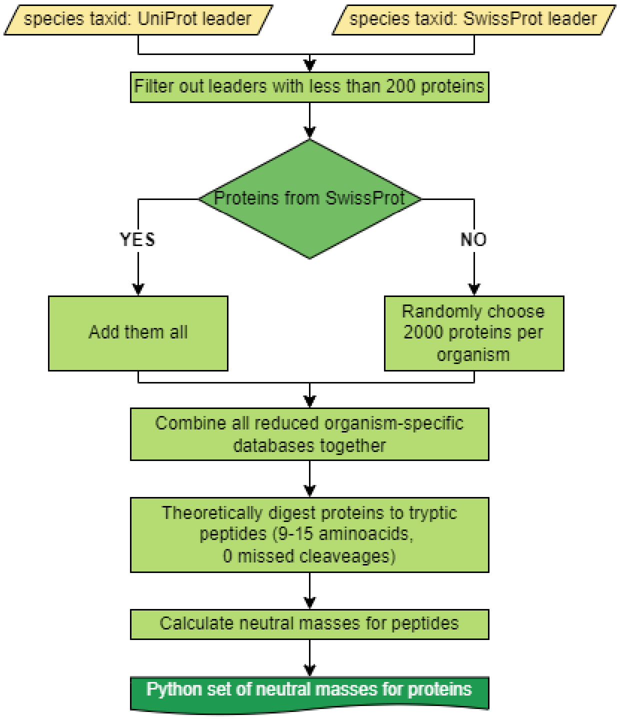
species taxid: UniProt leader
species taxid: SwissProt leader
Filter out leaders with less than 200 proteins
Proteins from SwissProt
YES
NO
Add them all
Randomly choose 2000 proteins per organism
Combine all reduced organism-specific databases together
Theoretically digest proteins to tryptic peptides (9-15 aminoacids, 0 missed cleaveages)
Calculate neutral masses for peptides
Python set of neutral masses for proteins

VII

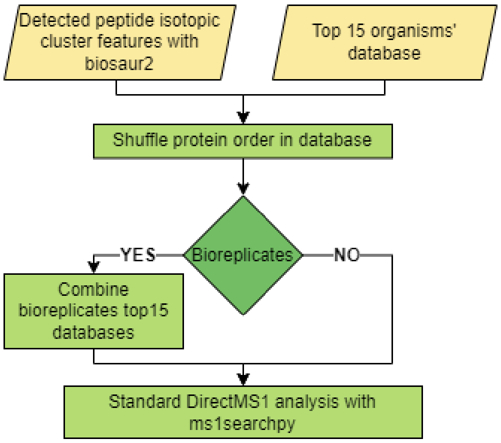
Detected peptide isotopic cluster features with biosaur2
Top 15 organisms' database
Shuffle protein order in database
Bioreplicates
YES
NO
Combine bioreplicates top15 databases
Standard DirectMS1 analysis with ms1searchpy

VI

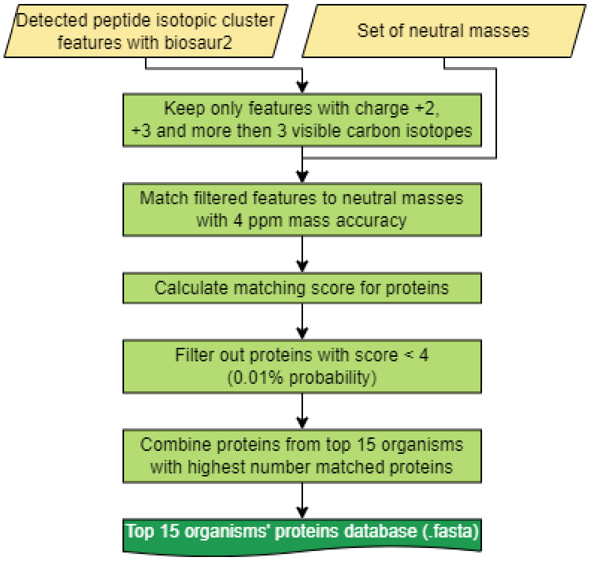
Detected peptide isotopic cluster features with biosaur2
Set of neutral masses
Keep only features with charge +2, +3 and more then 3 visible carbon isotopes
Match filtered features to neutral masses with 4 ppm mass accuracy
Calculate matching score for proteins
Filter out proteins with score < 4 (0.01% probability)
Combine proteins from top 15 organisms with highest number matched proteins
Top 15 organisms' proteins database (.fasta)

**Figure S2.** The normalized densities of $log_2FC$ distribution for ABRF2/ABRF3 comparison. The green line stands for the actual fold change calculated from the peptide masses used for spike preparation (peptide masses are written in strain subtitles) and the red line stands for the calculated ratio.

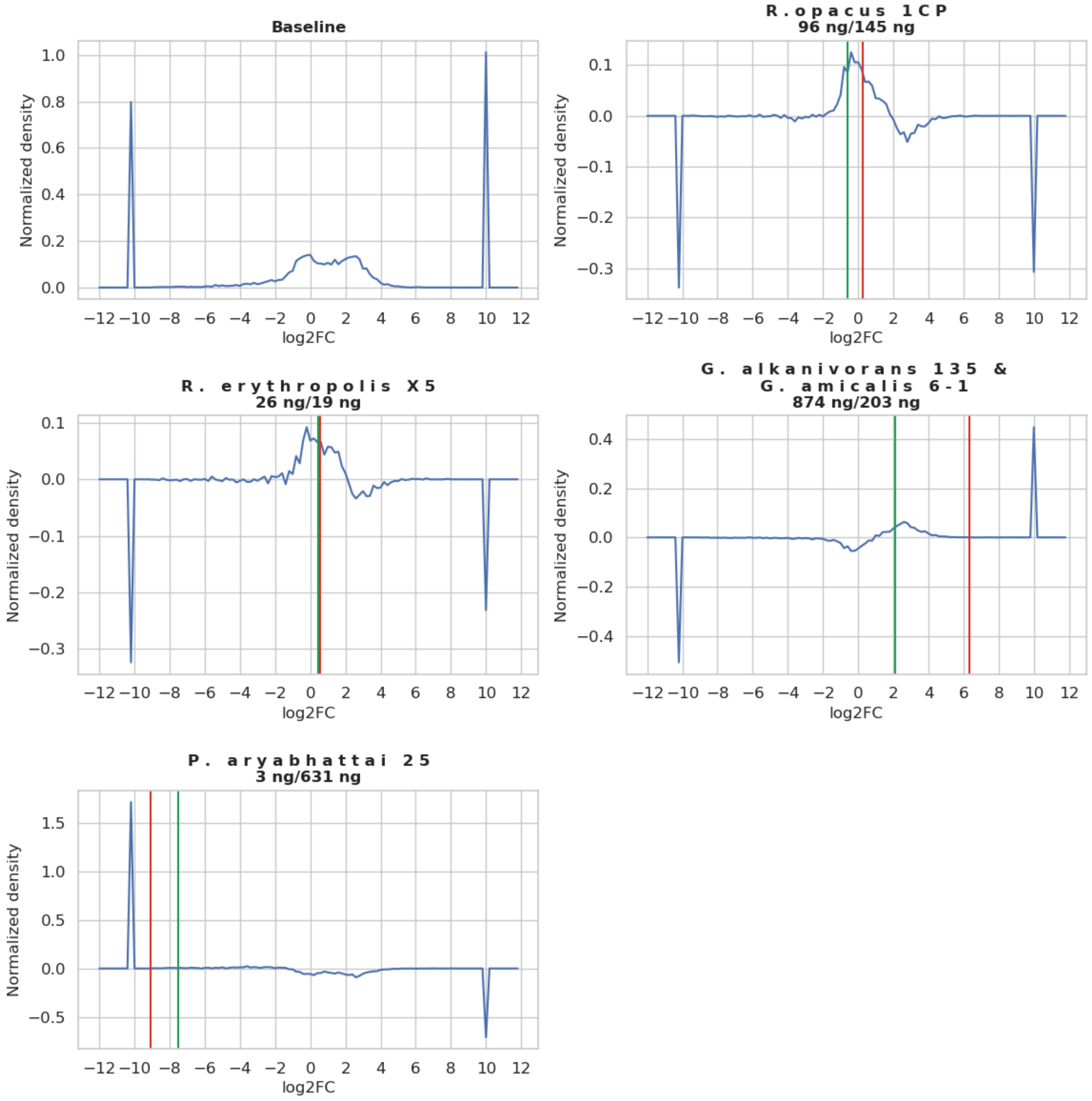

**Figure S3.** The normalized densities of $log_2FC$ distribution for M2.2/M2.1 comparison. The green line stands for the actuall value, calculated from peptide masses (shown as plots subtitles) and the red line stands for the calculated ratio.

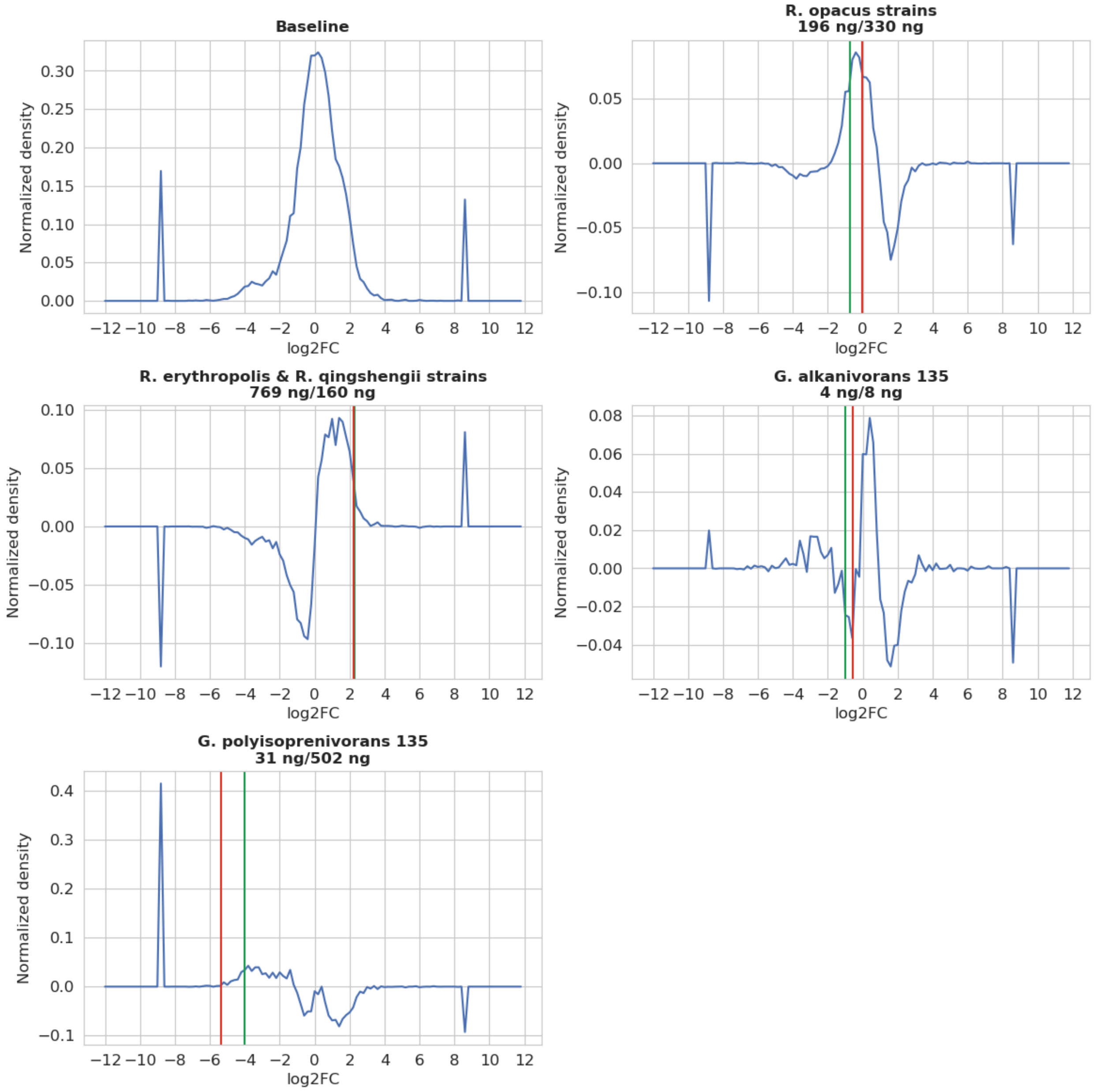

**Appendix B.** Protein signatures involved in degradation of *n*-alkanes at 6 $^{0}$C and at 28 $^{0}$C.

### *n*-Alkane degradation at 6 $^{0}$C:

*Oxidation to alcohols*:

QEX08599 cytochrome P450
QEX09356 alkane 1-monooxygenase
QEX11207 alkyl hydroperoxide reductase AhpD
QEX10447 3D-(3,5/4)-trihydroxycyclohexane-1,2-dione acylhydrolase (decyclizing) iolD

*Oxidation to aldehydes with alcohol dehydrogenases*

QEX08400 alcohol dehydrogenase catalytic domain-containing protein
QEX09399 aldo/keto reductase
QEX13597, QEX13509, QEX08596 SDR family oxidoreductase

*Oxidation to fatty acids, fatty acid metabolism*

QEX08594 fatty acid--CoA ligase
QEX08597 aldehyde dehydrogenase family protein
QEX14410 NAD-dependent succinate-semialdehyde dehydrogenase
QEX13735 3-hydroxyacyl-CoA dehydrogenase
QEX12285 long-chain fatty acid--CoA ligase
QEX12224 CoA-acylating methylmalonate-semialdehyde dehydrogenase
QEX13318 acyl-CoA synthetase
QEX09418 acetyl-CoA C-acetyltransferase
QEX11854 1-acyl-sn-glycerol-3-phosphate acyltransferase
QEX11472 NAD(P)/FAD-dependent oxidoreductase
QEX08857 thiolase family protein
QEX09951 N-acetyltransferase
QEX12147 type I polyketide synthase

*Exopolysaccharide production*

QEX09055 2-isopropylmalate synthase leuA
QEX10519 2-oxo-4-hydroxy-4-carboxy-5-ureidoimidazoline decarboxylase GN=uraD
QEX12651 glucose-6-phosphate isomerase
QEX11399 phosphoenolpyruvate carboxylase
QEX12072 glycosyl hydrolase
QEX11730 imidazole glycerol phosphate synthase subunit HisF hisF
QEX12952 allantoicase
QEX08969 MMPL family transporter

*Iron transport*

QEX13740 diiron oxygenase
QEX13741 4Fe-4S ferredoxin
QEX09379 ABC transporter substrate-binding protein
QEX09730 ABC transporter ATP-binding protein

### Chaperons and stress proteins increased at 6 $^{0}$C:

QEX09757 DNA starvation/stationary phase protection protein
QEX11651 universal stress protein
QEX11169 PspA/IM30 family protein (phage shock protein (psp) operon (pspABCDE))
QEX09859 ATP-dependent chaperone ClpB GN=clpB
QEX13151 molecular chaperone HtpG GN=htpG
QEX09823 molecular chaperone DnaK GN=dnaK
QEX09986 chaperonin GroEL groL

### 30S / 50S ribosomal proteins lost at 6 $^{0}$C:

bg|QEX10305.1|QEX10305.1_RHOER 50S ribosomal protein L29
bg|QEX10302.1|QEX10302.1_RHOER 50S ribosomal protein L22
bg|QEX10300.1|QEX10300.1_RHOER 50S ribosomal protein L2 GN=rplB

bg|QEX10312.1|QEX10312.1_RHOER 50S ribosomal protein L6
bg|QEX10313.1|QEX10313.1_RHOER 50S ribosomal protein L18
bg|QEX12550.1|QEX12550.1_RHOER 50S ribosomal protein L25/general stress protein Ctc
bg|QEX10338.1|QEX10338.1_RHOER 50S ribosomal protein L13 GN=rplM
bg|QEX13951.1|QEX13951.1_RHOER 50S ribosomal protein L9
bg|QEX10829.1|QEX10829.1_RHOER 30S ribosomal protein S16
bg|QEX10167.1|QEX10167.1_RHOER 50S ribosomal protein L11 GN=rplK
bg|QEX11657.1|QEX11657.1_RHOER 30S ribosomal protein S1
bg|QEX10296.1|QEX10296.1_RHOER 30S ribosomal protein S10 GN=rpsJ
bg|QEX10297.1|QEX10297.1_RHOER 50S ribosomal protein L3
bg|QEX13952.1|QEX13952.1_RHOER 30S ribosomal protein S18 GN=rpsR
bg|QEX10811.1|QEX10811.1_RHOER 50S ribosomal protein L28 GN=rpmB
bg|QEX10308.1|QEX10308.1_RHOER 50S ribosomal protein L24
bg|QEX12056.1|QEX12056.1_RHOER 50S ribosomal protein L21 GN=rplU
bg|QEX10168.1|QEX10168.1_RHOER 50S ribosomal protein L1
bg|QEX10171.1|QEX10171.1_RHOER 50S ribosomal protein L7/L12
bg|QEX10309.1|QEX10309.1_RHOER 50S ribosomal protein L5 GN=rplE
bg|QEX10170.1|QEX10170.1_RHOER 50S ribosomal protein L10

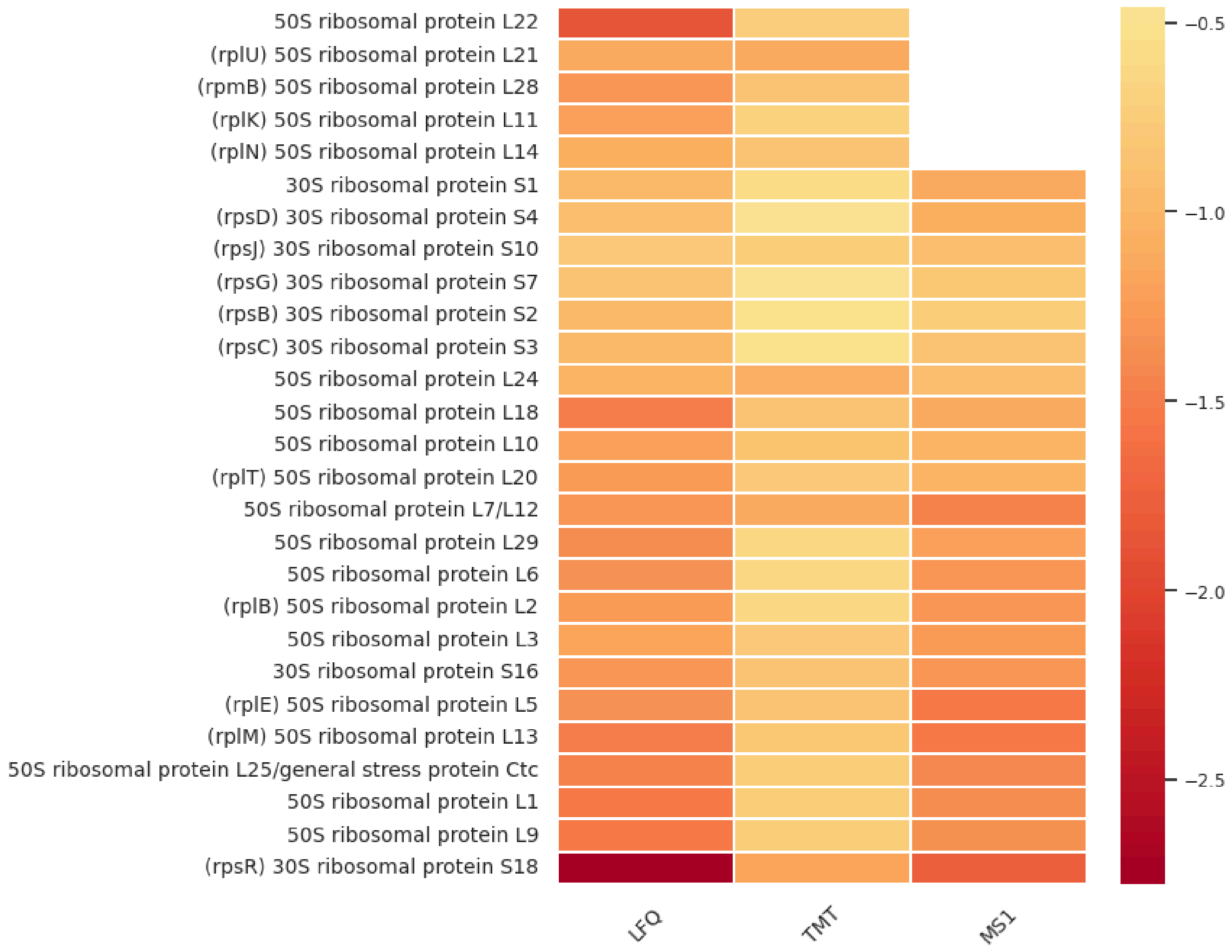


### ***n*-Alkane degradation at 24 $^0$C:**

*Oxidation to alcohols*:

QEX08601, QEX08447 cytochrome P450

*Oxidation to aldehydes:*

QEX10266 NDMA-dependent alcohol dehydrogenase
QEX10225 NDMA-dependent methanol dehydrogenase mdo
QEX10036 NAD(P)-dependent alcohol dehydrogenase

*Oxidation to fatty acids / fatty acid metabolism*:

QEX12404 long-chain-acyl-CoA synthetase
QEX10265 aldehyde dehydrogenase family protein
QEX10256 aldehyde dehydrogenase family protein
QEX08444 aldehyde dehydrogenase
QEX14300 acyl-CoA dehydrogenase
QEX13372 acyl-CoA dehydrogenase
QEX11983 3-oxoacyl-ACP reductase
QEX10534 acyl-CoA carboxylase subunit beta
QEX09026 3-hydroxyacyl-CoA dehydrogenase
QEX10443 CoA-acylating methylmalonate-semialdehyde dehydrogenase
QEX08752 succinate-semialdehyde dehydrogenase (NADP(+))
QEX09550 glutathione peroxidase
QEX09398 NADH:flavin oxidoreductase/NADH oxidase family protein
QEX10098 bifunctional uroporphyrinogen-III C-methyltransferase/uroporphyrinogen-III synthase

*Exopolysaccharide production*

QEX12160 glycosyltransferase family 1 protein
QEX13966 inositol-3-phosphate synthase
QEX14217 phosphomethylpyrimidine synthase ThiC GN=thiC
QEX12111 glutamine synthetase type III
QEX11539 methionine synthase GN=metH
QEX12609 UTP--glucose-1-phosphate uridylyltransferase
QEX10340 phosphoglucosamine mutase
QEX11604 pseudouridine synthase

*Iron transport*

QEX11646 excinuclease ABC subunit UvrA GN=uvrA
QEX12370 ABC transporter substrate-binding protein
QEX11669 ABC transporter substrate-binding protein
QEX14497 ABC transporter substrate-binding protein
QEX13072 ATP-binding cassette domain-containing protein

### DNA and RNA processing, regulation of transcription and translation at 24 $^0$C:

QEX14268 class 1b ribonucleoside-diphosphate reductase subunit alpha GN=nrdE
bg|QEX10996.1|QEX10996.1_RHOER ribosome recycling factor
bg|QEX11232.1|QEX11232.1_RHOER RNA polymerase sigma factor
bg|QEX12896.1|QEX12896.1_RHOER nuclear transport factor 2 family protein
bg|QEX10325.1|QEX10325.1_RHOER DNA-directed RNA polymerase subunit alpha
bg|QEX09649.1|QEX09649.1_RHOER GIY-YIG nuclease family protein
bg|QEX12425.1|QEX12425.1_RHOER translational GTPase TypA GN=typA
bg|QEX10183.1|QEX10183.1_RHOER DNA-directed RNA polymerase subunit beta GN=rpoB
bg|QEX14364.1|QEX14364.1_RHOER trigger factor
QEX10166 transcription termination/antitermination protein NusG GN=nusG
bg|QEX12054.1|QEX12054.1_RHOER GTPase ObgE GN=obgE
bg|QEX09300.1|QEX09300.1_RHOER CarD family transcriptional regulator
bg|QEX11602.1|QEX11602.1_RHOER ribosome biogenesis GTPase Der
bg|QEX11439.1|QEX11439.1_RHOER transcriptional regulator
bg|QEX12191.1|QEX12191.1_RHOER transcription termination factor Rho
bg|QEX10597.1|QEX10597.1_RHOER DEAD/DEAH box helicase

bg|QEX11824.1|QEX11824.1_RHOER cell division protein SepF
bg|QEX12264.1|QEX12264.1_RHOER TenA family transcriptional regulator
bg|QEX11827.1|QEX11827.1_RHOER cell division protein FtsZ GN=ftsZ
bg|QEX11178.1|QEX11178.1_RHOER recombinase RecA GN=recA
bg|QEX10184.1|QEX10184.1_RHOER DNA-directed RNA polymerase subunit beta'
bg|QEX10212.1|QEX10212.1_RHOER elongation factor G GN=fusA
bg|QEX08679.1|QEX08679.1_RHOER alpha/beta fold hydrolase
